# Clustering within a single-component biomolecular condensate

**DOI:** 10.1101/2025.08.18.670948

**Authors:** Ankith Sharma, Abhishek Sau, Sandeep Dave, Sumangal Roychowdhury, Sebastian Schnorrenberg, Rajdeep Chowdhury, Saskia Hutten, Dorothee Dormann, Siegfried M. Musser

**Author notes:** Equal Contributions (Ankith Sharma, Abhishek Sau).

## Abstract

Biomolecular condensates (BMCs) are assemblies of hundreds to many thousands of macromolecules within cells that are organized without physical barriers. Condensate function is dictated not only by its molecular composition, but also by substructural organization and molecular mobility. One hypothesis for the onset of multiple protein aggregation diseases is that the increased densities of specific proteins within BMCs promotes the formation of solid inclusions. However, deciphering the internal structural and functional properties of BMCs at the nanoscale and identifying the initiating events of inclusion formation requires tools with high spatiotemporal precision. Here we show using single molecule and other microscopy approaches that single component Fused in Sarcoma (FUS) condensates exhibit confinement and contain clusters with higher FUS density even at early timepoints. Upon aging, condensates displayed altered physical properties and reduced monomer mobility, and yet most FUS monomers diffused throughout the condensate within seconds. While an increase in connectivity over time explains reduced mobility, the large fraction of molecules retaining high mobility suggests a sponge-like structure rather than a system-spanning network. These findings indicate that a pseudo-equilibrium between distinct structural connectivities can exist within simple condensates, suggesting the potential for substantial structural and functional complexity of BMCs at the nanoscale.

---

Classical organelles are cellular compartments surrounded by phospholipid bilayer membranes. BMCs, such as stress granules, Cajal bodies, and nucleoli, constitute an additional organizational component of cells without a physical barrier to regulate exchange between compartments^1,2^. BMCs form as the result of a density transition with respect to one or more molecularly distinct species wherein it is more favorable for an assembly of macromolecules to coalsce rather than remain dispersed throughout the accessible volume^3,4^. Such assemblies (condensates) have a wide range of biological activities, including serving as biochemical factories and playing critical roles in sensing, activation, inactivation, nucleation and storage^1,5^. BMC self-assembly is driven by multivalency, often through numerous weak interactions between RNAs and intrinsically disordered regions of ‘scaffold’ proteins, which initiate and maintain condensate structure while simultaneously allowing access to ‘clients’ needed for functional activities^6–8^. In vitro, it is well-established that numerous proteins will individually phase separate to yield liquid-like droplets above a saturation concentration (*c*_sat_), which is dependent on physicochemical conditions^9–11^. The condensation process is termed liquid-liquid phase separation (LLPS) when the initial condensate has liquid-like properties, but alternate condensed states, such as a percolated network or a solid, are also possible either over time or coupled to the initial density transition^3,12,13^. Multiple phases within multi-component condensates exhibit a wide range of organizational structures, indicating that multiple factors influence miscibility^14,15^. Clusters or alternate assemblies can form below *c*_sat_, indicating complexity beyond simple Flory-Huggins models of phase separation^16–18^ and also revealing the potential for sub-diffraction-sized assemblies of phase separating proteins. The conditions under which distinct assemblies form and their interactions with and within condensates are not well-established. As typical physiological condensates are of micrometer size^7^, determining sub-structural and dynamical properties remains challenging. Nonetheless, clusters and amyloid fibrils within condensates have been identified^19–23^.

Single-molecule super-resolution approaches are in principle ideally suited to probe BMCs; however, data collection with traditional fluorophores is often limited by two characteristics of BMCs, their small size and the limited or slow exchange with the bulk. Single molecule fluorescence (SMF) measurements are pointillistic, i.e., they rely on detecting a single fluorophore with a light emission pattern typically multiple hundreds of nanometers in diameter to construct an image or characterize behaviors^24,25^. For micrometer sized BMCs, single molecule detection generally requires that at most a few fluorophores within a condensate are detectable at any given moment. In addition, condensates are generally highly viscous^26–28^, and thus the molecular exchange required to visualize subsequent molecules after photobleaching is inherently slow, particularly after aging when molecular mobility is frequently substantially reduced^28,29^. Here, we used spontaneously blinking and caged fluorophores to measure translational and rotational mobility and to identify differences in molecular densities within Fused in Sarcoma (FUS) condensates. FUS has been identified as the major protein in the inclusions (aggregates) found in some patients with amyotrophic lateral sclerosis (ALS) and frontotemporal dementia (FTD)^29,30^. Using three-dimensional (3D) imaging, we identified stable cluster-like microenvironments far from the coverslip surface. The immobility of such clusters within a medium dominated by higher mobility suggests an assembly with variable connectivity, i.e., a relatively open sponge-like structure rather than a system spanning percolated network.

## FUS condensates contain clusters

Condensates were formed from wild-type full-length FUS purified from bacterial cells (**Fig. 1a,b** and **Extended Data Fig. 1**). For visualization/interrogation of condensates, a dye-labeled A2C mutant (FUS^A2C^; **Fig. 1a**) was added at ≤ ∼0.14% of the total FUS protein; the native cysteines within the zinc finger domain were protected against maleimide dye labeling (**Extended Data Fig. 1**), presumably due to the C4-type chelation of Zn^2+[31]^. Previous work indicated that FUS distributes heterogeneously within initially liquid-like condensates and that inhomogeneities increase as condensates age^32,33^. To explore such structural changes, we used super-resolution microscopy to assess how the FUS density distribution evolves over time. We took advantage of the spontaneously blinking dye, JF635b (**Fig. 1c**), with an on/off ratio (duty cycle) of ∼0.05%, which allowed for the individual detection of single fluorophores within a crowded environment without the need for external reagents to induce blinking^34^ or exchange/replenishment from the bulk. Using FUS condensates spiked with 0.14% FUS^A2C^-JF635b, fluorophores were localized in 3D via astigmatism imaging^35,36^ (**Fig. 1d-f**, **Supplementary Video 1** and **Extended Data Fig. 2**), yielding an average of ∼2831 localizations/µm^3^ **(Extended Data Fig. 3**). Variable localization densities (PALM intensities) were observed throughout the condensates with an apparent increase in higher density regions at longer aging times (**Extended Data Fig. 2c**). A random distribution of dye-labeled proteins throughout a homogeneous condensate, however, will naturally lead to variable measured densities, which need to be calibrated to the total density within the volume. To evaluate whether observed densities were higher than would be expected from stochasticity, we simulated random distributions (**Fig. 1h,j,l**) with average overall densities identical to those experimentally observed (**Fig. 1g,i,k**). Density histograms revealed a wider range for the experimental data (**Extended Data Fig. 2e**), verifying that the observed higher density regions (clusters) were bona fide physical features of the condensates. In some particularly dense regions at longer aging times (e.g., 7 days), irregular localization distributions were observed consistent with assemblies of clusters or filamentous structures (**Fig. 1m**). The number of localizations above the high-density simulation threshold increased with aging time (**Fig. 1n**). Well-separated clusters had an average diameter of ∼150 nm (**Fig. 1o,p**), a value that is likely broadened by slow movement over the imaging time. Here, the term ‘clusters’ is a functional definition, denoting regions within 3D space where a higher density of dye localizations were obtained. Whether this occurred because of molecular density differences or environmental differences that influenced blinking frequency – or a combination of these effects – has not been established by these data. Nonetheless, the higher densities of dye localizations identified regions of distinct physicochemical properties with measurable shapes (**Fig. 1m,o,p**).

**Fig. 1.**
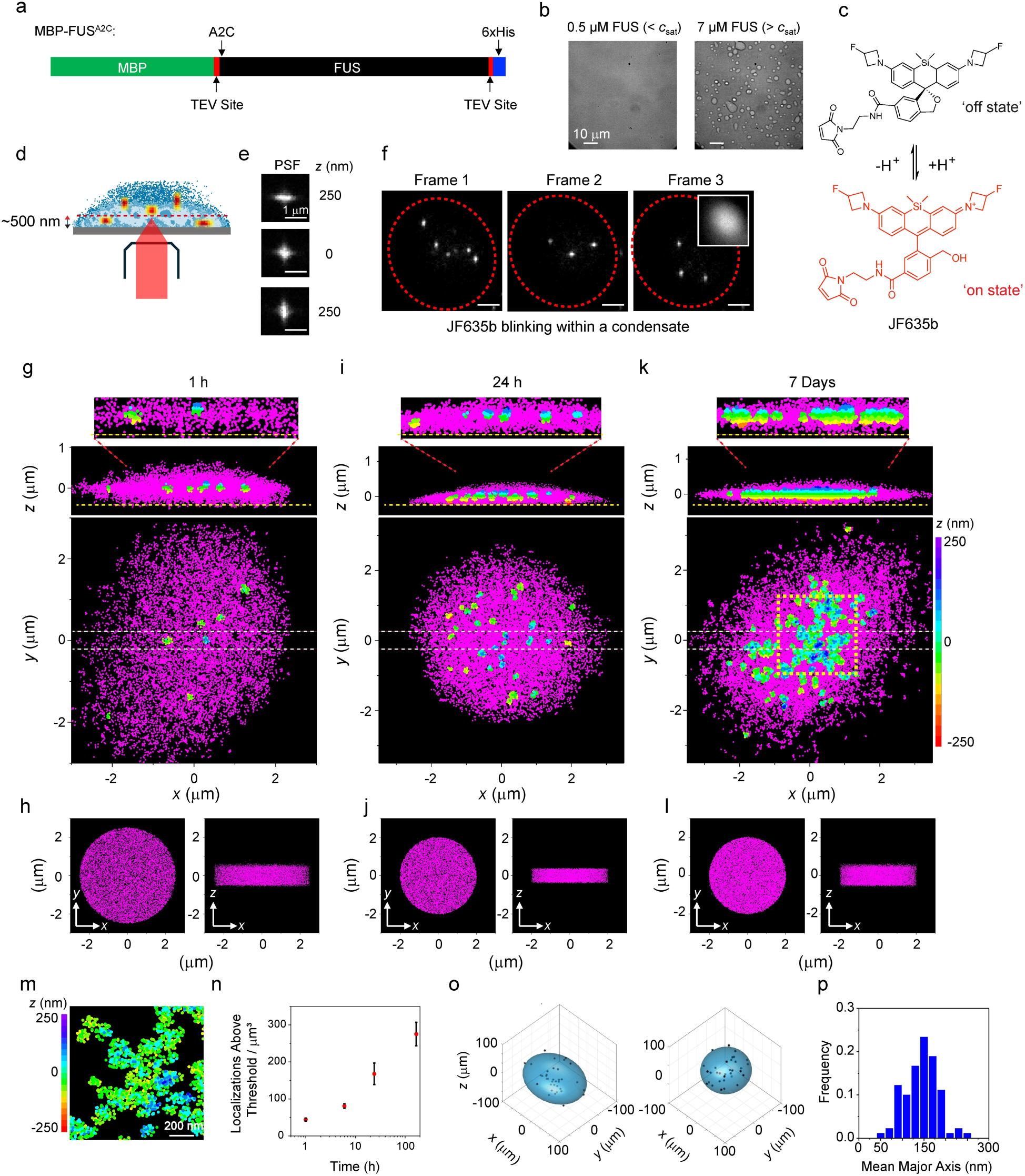
Clustering within FUS condensates observed in localization density maps. **a,b,** Condensates formed from full-length FUS. The full-length wild-type FUS protein as well as the A2C mutant were purified with an N-terminal maltose binding protein (MBP) solubility tag and C-terminal 6xHis tag (**a**). Condensates formed above *c*_sat_ when the two tags were released by TEV protease activity (**b**, **Extended Data Fig. 1**). All condensates in this paper were performed at a final FUS concentration of 7 µM. **c-f**, Localization of single fluorophores in three dimensions (3D) at high dye densities. MBP-FUS^A2C^ was labeled with the self-blinking dye JF635b (**c**). With a focal plane ∼500 nm above the coverslip (**d**), astigmatism imaging was used to locate single JF635b fluorophores within 3D – the *xy*-coordinates were determined from the centroid and the *z*-coordinate was determined from the extent of spot elongation in *x* or *y* (**e**, see Methods). Due to the spontaneous blinking of JF635b, the molecules observed in subsequent image frames change rapidly (**f**, **Supplementary Video 1**). The inset in frame 3 of **f** is the average of the fluorescence observed over 30,363 frames, identifying the location of the condensate (red dashed ovals). Scale bars (**f**) = 2 µm. **g-l,** Increased clustering with longer aging times. 3D localization density maps were obtained at three aging times (**g**,**i**,**k**) using FUS condensates containing ∼0.14% of FUS^A2C^-JF635b and imaged as in **f** for ∼20 min at 30 ms/frame. Scatter plots of all localizations (average of 2831 localizations/µm^3^) are shown. To confirm that the observed experimental distributions (**g**,**i**,**k**) were non-random, simulations of random distributions within a cylinder (**h**,**j**,**l**) were generated with densities matching those in the experimental condensates. Regions of high density within the condensates and simulations were identified by DBSCAN (**Extended Data Fig. 2** and Methods). Localizations within identified clusters are colored by *z*-scale amongst all localizations (*magenta*). Expanded views (top of **g**,**i**,**k**) indicate that the cluster regions are found within the condensate and not stuck to the coverslip surface. Dashed lines in the *xz* images indicate the position of the coverslip. Additional datasets in **Extended Data Fig. 3**. **m,** Interconnected clusters. Magnified region from the yellow boxed region of the condensate in **k**. **n**, Increased clustering over time. The number of localizations within high-density regions increased as the condensates were aged from 1 h to 7 days (*n* = 3 for all timepoints; error bars are standard deviations). See **Extended Data Fig. 2f** for an explanation of the threshold. **o,p,** Cluster size. Individual clusters (1-24 h) were fit to an ellipsoid (**o**). Clusters had a mean major axis length of ∼150 nm (**p**, *n* = 90 clusters).

The observed clustering requires immobility of the cluster over the observation window. Thus, since data were collected over 20 min, the higher densities observed in the localization maps of **Fig. 1g,i,k** are consistent with entities that have extreme immobility, although molecules could in principle move into and out of the clusters during the imaging time. Due to the known maturation of FUS condensates to more immobile phase states^37–39^, the immobility of clusters after days to a week of aging is unsurprising. Unexpectedly, however, some clusters were already observable at the 1 h timepoint (**Fig. 1g**), indicating that such clusters were immobilized already at a very early step in the aging process. Due to the 3D imaging strategy, these clusters were clearly detected within the condensate interior and therefore they were not stuck to the coverslip surface (**Fig. 1g**, top). This indicates the presence of a high viscosity medium or an internal structure that restricts cluster movement shortly after condensate formation.

## FUS mobility decreases over time

To assess the presence and mobility of clusters at earlier timepoints, we employed a faster imaging method with a low concentration of a normal (non-blinking) dye. Using wide-field microscopy, slow- and fast-moving species of FUS^A2C^-JF549 were visualized within condensates, consistent with a reduced mobility of some of the dye-labeled protein molecules due to their presence in clusters. This was most readily observed after an initial short period (∼5 s) of photobleaching that occurred while searching and focusing (**Supplementary Videos 2**-**4**). The overall dynamics indicated liquid-like condensates at early time points (e.g., 1 h) – condensate fusion was observed, and most fluorescent molecules moved rapidly (**Supplementary Video 2**). However, fluorophore immobility increased at later timepoints (**Supplementary Videos 3-4**). By comparing images with fixed delay times, the distribution of the dye-labeled protein fluctuated rapidly after 1 h of aging, indicating high mobility, but mobility decreased over a week of aging (**Fig. 2a,b**). In agreement with previous work^16^, small assemblies of FUS (≤ 200 nm diameter) formed at low concentration, as assayed by dynamic light scattering (**Fig. 2c** and **Extended Data Fig. 4a**,b). However, whether such assemblies reflect a distinct physical state forming below *c*_sat_ (such as the nanoclusters^19,21^ or pre-percolation clusters^3,20^observed by others), or simply oligomers or small condensates remains uncertain. The intensities of slowly moving spots within the condensates largely corresponded to that of a single dye molecule, and the movement behavior were consistent with these spots corresponding to assemblies ∼100-200 nm in diameter (see **Extended Data Fig. 4d-f**). Such large single dye-containing assemblies are unlikely to have been induced by the hydrophobic nature of a single dye molecule. These observations are consistent with a model in which clusters that initially formed below *c*_sat_ during the rapid increase in the concentration of free FUS by TEV protease activity (see Methods, **Fig. 1a,b** and **Extended Data Fig. 1**) were sometimes trapped within growing condensates. However, these data do not rule out the possibility that intra-condensate clusters sometimes or exclusively assembled after condensate formation, and that intra-condensate clusters are physicochemically distinct from extra-condensate assemblies.

**Fig. 2.**
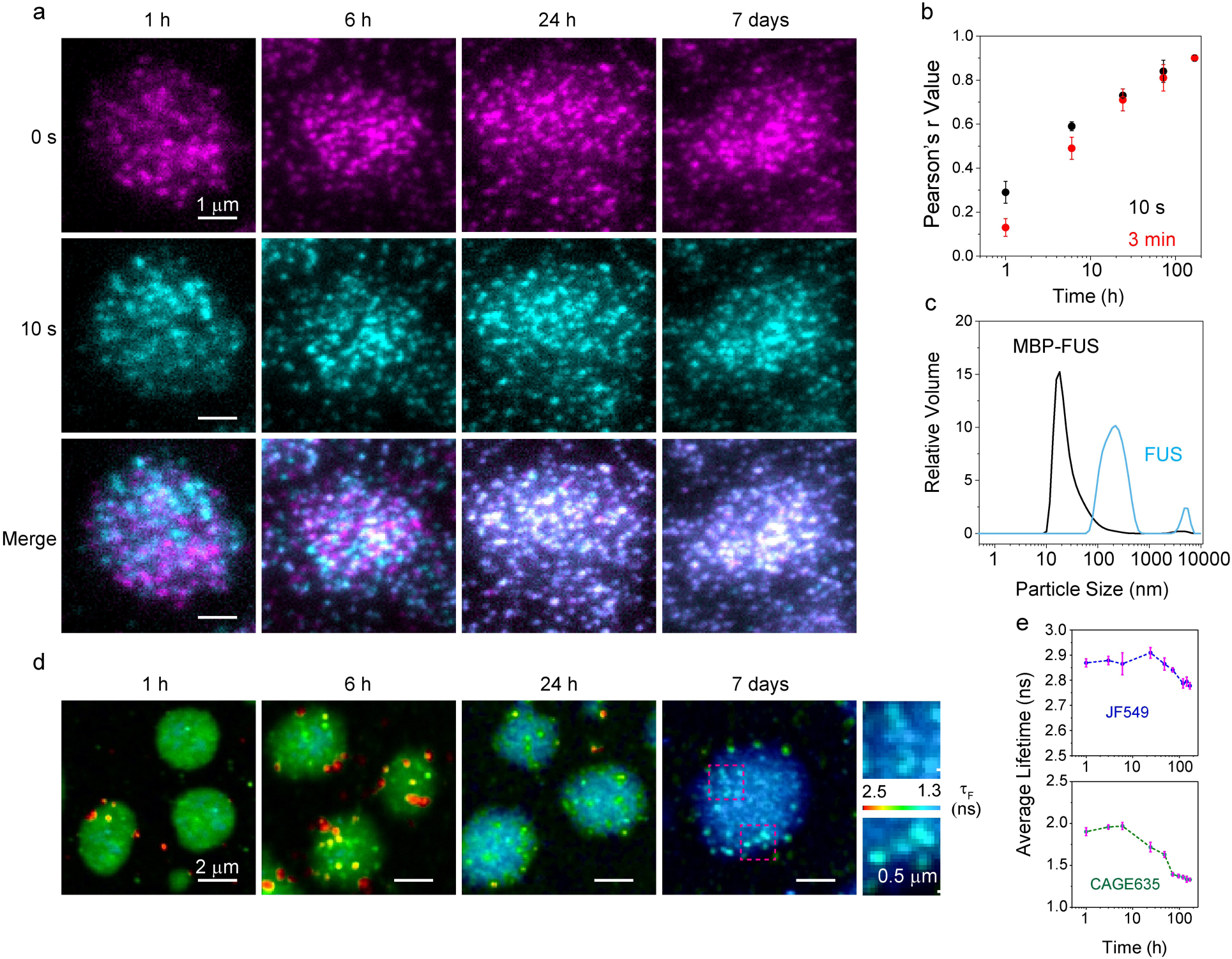
Condensate heterogeneity assessed by dye mobility and fluorescence lifetime. **a,b,** Reduced FUS mobility upon condensate aging. FUS condensates with ∼0.5 nM of FUS^A2C^-JF549 were imaged by epifluorescence after different aging timepoints, as indicated (**a**). At every aging timepoint, images were obtained at *t* = 0 s, 10 s and 180 s and compared. To visually illustrate mobility (**a**), *t* = 0 s (top, *magenta*) and *t* = 10 s (middle, *cyan*) frames were merged (bottom), where immobility is indicated as *blue* to *white*, depending on overall intensity. Fluorescence colocalization after the imaging delay was quantified using Pearson’s overlap coefficient (**b**; *n* = 3 condensates for all timepoints). **c,** Nanoscale FUS assemblies at low concentration. Dynamic light scattering (DLS) indicates that assemblies of ∼200 nm diameter were formed at 0.5 µM FUS and 75 mM NaCl (see **Extended Data Fig. 4a** for the effect of salt concentration). **d,** FLIM of FUS condensates containing FUS^A2C^-CAGE635 (∼10 nM). The color code represents the amplitude-weighted average fluorescence lifetime per pixel. Puncta are consistent with the clusters identified in Fig. 1. Enlarged images from the red dashed boxes of the 7 days image are shown on the right. These images reveal puncta clustering similar to the cluster of clusters observed in Fig. 1m. **e,** Average fluorescence lifetime at different condensate aging times for the JF549 and CAGE635 dyes. Plotted are the weighted average lifetimes from a double-exponential fit to combined data for each condensate (*n* = 10 condensates for all timepoints). See **Extended Data Fig. 5** for representative data at all time points.

## Clusters are internally distinct

Fluorescence lifetime imaging microscopy (FLIM) was used to test whether there are detectable physical differences in the fluorophore environments within the bulk and cluster regions. Fluorescence lifetime (1_F_) is sensitive to pH, solvent polarity, viscosity, and the presence of quenchers^40,41^. FLIM images were collected within 3 min using a scanning confocal imaging scheme. Puncta were readily identifiable in the condensates, particularly at longer timepoints, suggesting that the fluorophores in clusters have distinct lifetimes from the fluorophores in the bulk (**Fig. 2d**). A longer average 1_F_ for both JF549 (a continuously ‘on’ dye) and CAGE635 (a photoactivatable dye) was observed for puncta, although the Δ1_F_ between the bulk and puncta was larger for CAGE635 (**Fig. 2d** and **Extended Data Fig. 5**). The weighted average 1_F_ for CAGE635 decreased with condensate age, consistent with a decreased pH and/or solvent polarity, but the magnitude of the decrease was larger for CAGE635, indicating a differential sensitivity of the dyes to the condensate environment (**Fig. 2e**). Assuming that the puncta observed in FLIM images are indeed the clusters observed in the density maps (**Fig. 1g,i,k,m**), these data support the conclusion from the density maps that the internal environment of the clusters and the bulk of the condensate are distinct physical environments. After 7 days, the puncta detected in FLIM images are more crowded and seemingly merge to form extended structures (**Fig. 2d**), similar to what was observed earlier in the density maps (**Fig. 1m**), although it must be noted that the FLIM images are lower resolution and 2D (not 3D). The higher 1_F_ for the puncta identified in FLIM images is consistent with an increased viscosity^41^, which is expected for a higher FUS density in clusters (**Fig. 1**). However, changes in local pH and solvent polarity cannot be ruled out.

## Three diffusional mobilities

While some FUS molecules exhibited extreme immobility (**Fig. 2a,b** and **Supplementary Videos 2**-**4**), this represented a small fraction of the total FUS population. We therefore sought to assess the number and fractional composition of distinct mobilities. While the JF635b blinking dye allowed for the individual localization of tens of thousands of single fluorophores per condensate, it switched off too rapidly to obtain the reasonable-length single particle trajectories necessary to adequately characterize translational mobility. We therefore switched to CAGE635, a UV photoactivatable dye, to obtain longer tracks from a series of localizations. Condensates were imaged at 5 ms/frame using astigmatism imaging for ∼20 minutes, yielding an average of 806 tracks/condensate with a minimum of 5 localizations/track (**Fig. 3a**). Two distinct behaviors were immediately discernable, namely, tracks that explored a significant fraction of the condensate and tracks that exhibited highly localized movements, consistent with confinement to clusters (**Fig. 3a**). Jump histograms of trajectories confined to clusters yielded the diffusion coefficient *D*_1_ ≈ 0.01 µm^2^/s for all timepoints (**Fig. 3b** and **Extended Data Fig. 6**). As these jump steps were comparable in magnitude to the localization precision of the measurements, a simulation model was necessary to account for this localization error to accurately estimate *D* (**Fig. 3b**). Two additional diffusion coefficients (*D*_2_ and *D*_3_) were needed to fit a jump step histogram comprised of all the tracking data at a given timepoint (**Fig. 3c-f** and **Extended Data Fig. 6d-f**). The values of *D*_1_, *D*_2_, and *D*_3_ were largely unchanged with aging time (**Fig. 3d**; 0.014, 0.26, and 0.73 µm^2^/s, respectively, averaged over all timepoints); however, the fraction of *D*_3_ decreased (0.67 to 0.17), that of *D*_2_ increased (0.22 to 0.69), and that of *D*_1_ was fairly constant (∼0.08-0.14) over 7 days (**Fig. 3e**). Consequently, aging resulted in a reduction of the weighted average *D* by ∼40% (**Fig. 3g**). The diffusion data do not support a model in which the fraction of molecules within clusters increased substantially over time. Instead, the density maps (**Fig. 1g,i,k**) and narrow-field particle mobility data (**Fig. 2a,b** and **Supplementary Videos 2**-**4**) suggest that an increased immobility of clusters at longer aging times led to a higher fraction of cluster-confined molecules being interrogated within ‘static’ clusters. Accordingly, considering that cluster movement was likely significantly slower than FUS monomer movement, the movement of cluster-confined FUS can be expected to be largely the same whether the cluster was moving or static.

**Fig. 3.**
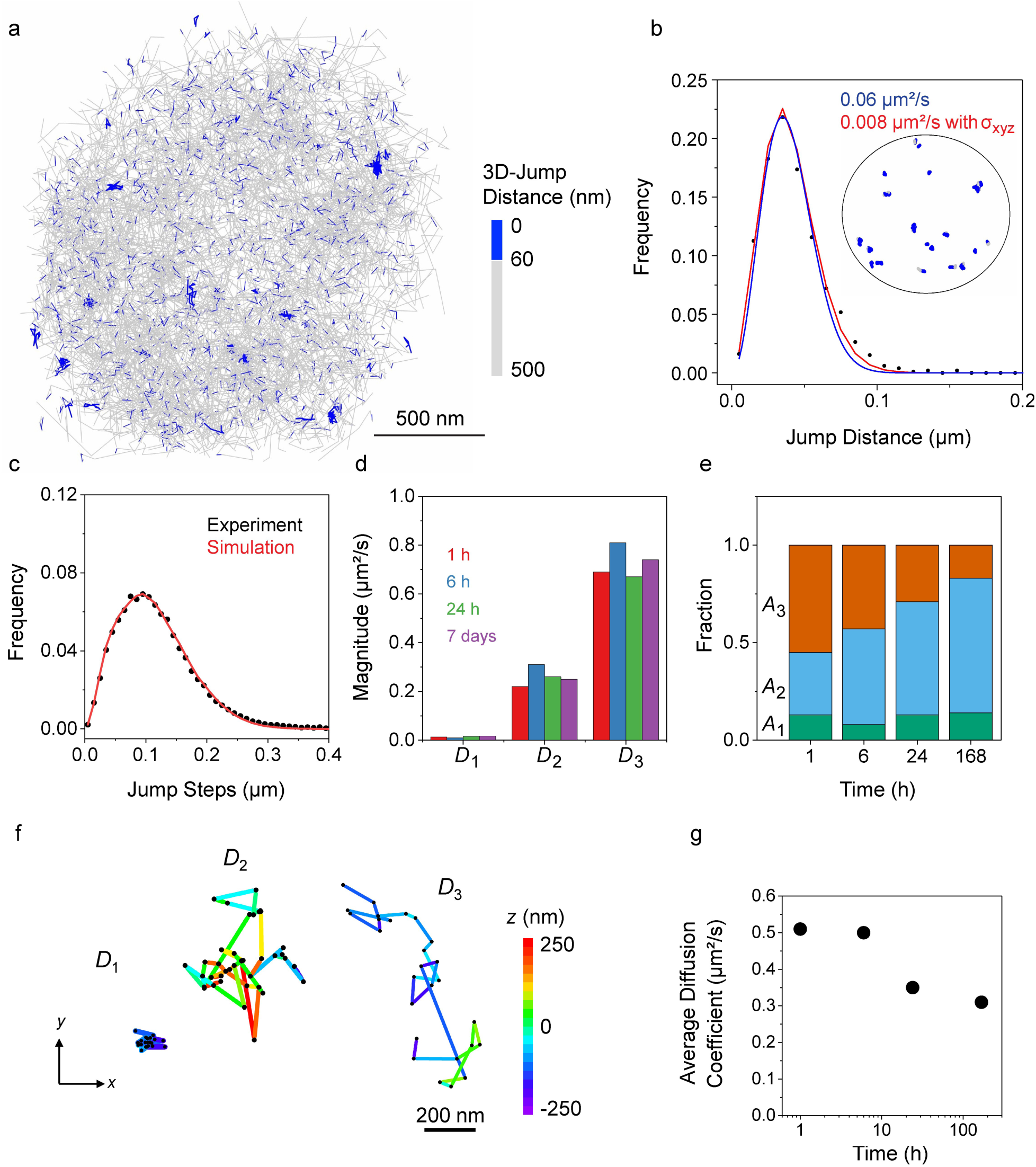
3D tracking reveals three distinct diffusive species of FUS within condensates. **a,** 3D diffusion behavior of FUS^A2C^-CAGE635 (1 nM) within a FUS condensate aged for 6 h. All 1227 trajectories with ≥ 5 localizations are shown. Highlighting the short (< 60 nm) jump steps (*blue*) identified confined trajectories. Only localizations with *z* = 0 ± 250 nm and ≥ 500 photons/localization were analyzed further. **b,** Jump histogram for confined trajectories, defined as those with an average jump step < 50 nm. As confirmed by the inset (from **a**), this selection identified trajectories localized to a small region. Disregarding measurement error, the fit (*blue*; **Eq. 1**) yielded a diffusion coefficient of 0.06 µm^2^/s. Accounting for localization precision (average values: σ_x_ = 12.7 nm, σ_y_ = 15 nm, and σ_z_ = 21 nm; see Methods and **Extended Data Fig. 6**), the diffusion coefficient was estimated as 0.008 µm^2^/s (*red* fit), thus illustrating that accounting for localization precision is essential for estimating slow diffusion coefficients. **c,** Jump step histogram after 6 h of aging (*n* = 10,682 jump steps, 1 condensate). Accounting for localization precision, three diffusion coefficients (0.005, 0.23, and 0.63 µm^2^/s for the data shown) were required to fit the data (see **Extended Data Fig. 6** for other fits). **d-g,** Diffusion parameters for different aging times. The three diffusion coefficients (**d**), their corresponding fractional contributions (**e**), selected tracks approximating the diffusion behavior of the three diffusion coefficients (**f**), and the weighted average diffusion coefficient (**g**) calculated from the values in **d** and **e** are provided (*n* = 3 condensates for all timepoints).

## Dynamics within clusters

The density maps indicated that clusters had dimensions below the diffraction limit (**Fig. 1o,p**) and the tracking data indicated highly localized movements of FUS monomers within clusters (**Fig. 3**). To further examine these issues with high spatiotemporal precision, 3D MINFLUX^42^ was used to characterize the diffusion behavior of slow-moving FUS proteins. MINFLUX is extremely photon efficient compared to diffraction-limited approaches, requiring ∼10-fold less photons to achieve a similar spatiotemporal localization precision for single fluorophores, which thus yields substantially longer particle trajectories^42–44^. A sufficiently long trajectory for a molecule diffusing within a cluster could, in principle, report on the estimated size and shape of the cluster. Unfortunately, due to the donut-shaped excitation pattern in MINFLUX that amplifies background over the signal from the fluorophore of interest, such measurements are particularly sensitive to background noise. FUS condensates exhibited a small but measurable fluorescence background that increased over time, which therefore limited us to early timepoint measurements and generally eliminated trajectories with higher particle mobility. This fluorescence background is at most a few percent of the total signal measured for the other fluorescence techniques reported in this paper, and hence, has a minimal influence on the other data reported here.

Trajectories obtained by MINFLUX for FUS within condensates aged for 3-6 h were obtained throughout the condensates (**Fig. 4a**). After quality control filtering (**Extended Data Fig. 7**), multiple trajectory behaviors were observed within 87 total tracks (**Fig. 4b** and **Extended Data Fig. 8**). Surprisingly, though these could be classified as ‘confined’, ‘partially confined’ and ‘unconfined’ trajectories, they yielded very similar jump step histograms (**Fig. 4c**), which could be fit by assuming two diffusion coefficients, 0.05 µm^2^/s and 0.008 µm^2^/s, with different levels of confinement (**Extended Data Fig. 7f,g**). These parameters also yielded simulated trajectories that closely resembled the experimental data (**Fig. 4b**). The average diffusion coefficient was ∼0.019 µm^2^/s, which we take, within experimental error, to reflect the *D*_1_ obtained by astigmatism imaging (**Fig. 3**). Mean-squared displacement curves yielded an estimated *D* = 0.01 µm^2^/s, also consistent with the previously determined *D*_1_, and confirmed that the tracked molecules experienced different levels of confinement (**Fig. 4d** and **Extended Data Fig. 7e**). Since the diffusion coefficients characterizing the different trajectories appear similar yet the trajectory behaviors appear distinct (**Fig. 4b-d** and **Extended Data Fig. 7d-g**), a reasonable explanation is that the environments of the diffusing molecules must therefore have been distinct, e.g., due to diffusion within clusters of different size and shape or due to a porous (semi-)static structure within the clusters or condensates. Some of the shorter trajectories (**Extended Data Fig. 8**) revealed an extreme confinement, which could reflect their embedment within a static scaffold.

**Fig. 4.**
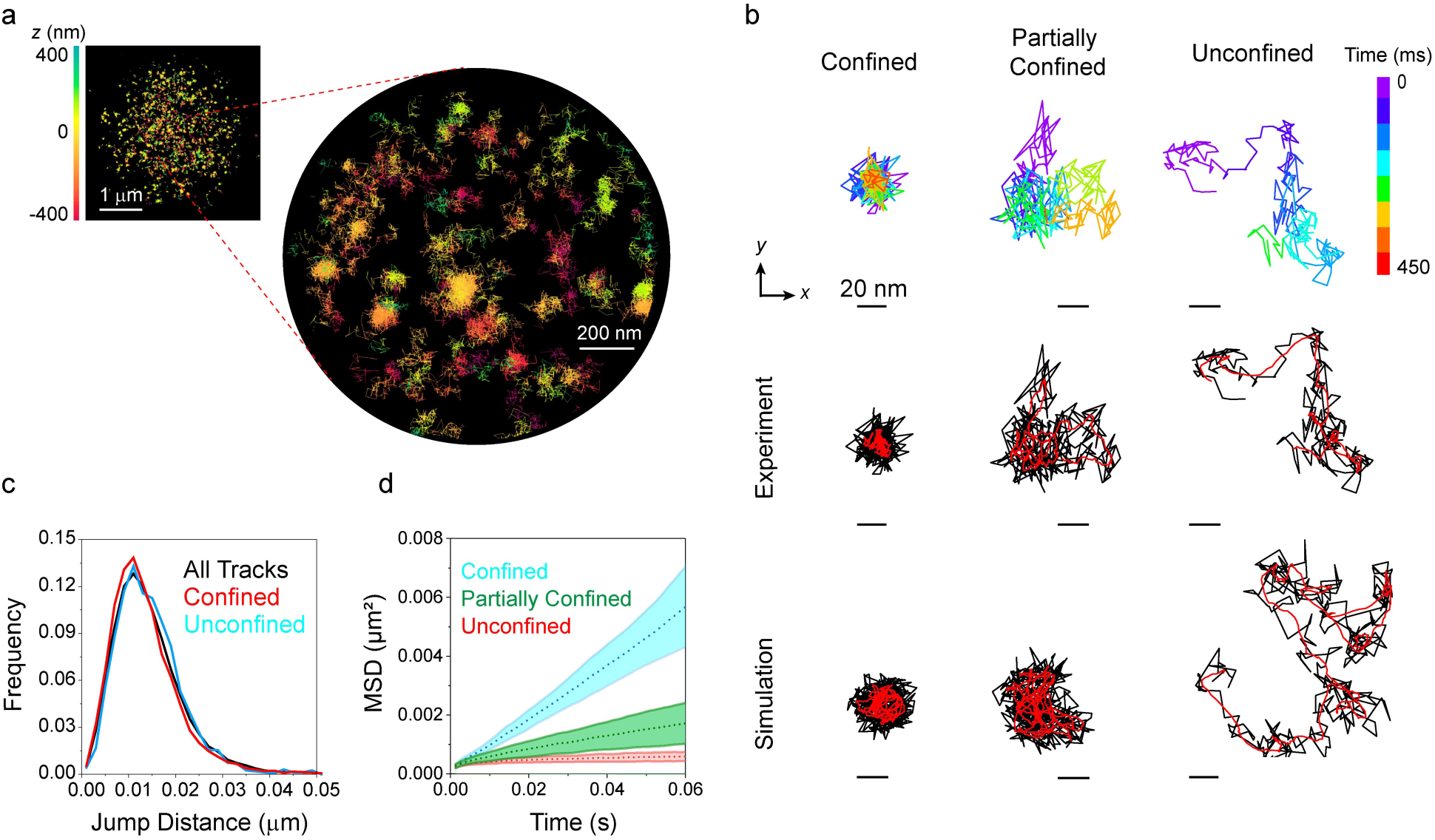
MINFLUX imaging reveals confinement. **a,** MINFLUX tracks of FUS^A2C^-CAGE635 (1 nM) obtained within a condensate aged for 3 h and imaged for 2 h. These are the raw unfiltered tracks with an average trajectory length of 150 localizations colored by *z* position. **b-d,** Variable confinement. A total of 87 trajectories from 4 condensates survived filtering (see Methods and **Extended Data Figs. 7 and 8**), and these exhibited different levels of confinement (**b**). Jump histograms calculated from confined, unconfined and all tracks were very similar (**c**), indicating that the diffusive motion was almost identical in all cases, with the exception of barriers to diffusion that caused the observed confinement. Based on mean squared displacement (MSD) curves (**d**), these tracks could be generally classified as confined (α = 0.44), partially confined (α = 0.79) or unconfined (α = 1.09), although a range of α ≈ 0.1-1.1 was observed for the individual trajectories (**Extended Data Fig. 7d,e**). Simulations indicated two diffusion coefficients [0.008 µm^2^/s (75%) and 0.05 µm^2^/s (25%)] with an average *D* = 0.019 µm^2^/s (**Extended Data Fig. 7f,g**). These values were used to generate simulated trajectories with different levels of confinement (**b**, *bottom row*), which differ in the spherical volume the particle was allowed to move in (radius = 25-1000 nm). The trajectories (*black*) in **b** are shown with a moving centroid calculated from 10 localizations (*red*). The experimentally observed MINFLUX tracks (**b**, *middle row*) are also colored based on time (**b**, *top row*).

## Extreme dye immobility

FUS condensate aging favors the formation of solid assemblies^37–39^. Since we observed no evidence of fibril formation, we therefore used single molecule rotational diffusion (SiMRoD) microscopy^45^ to explore whether microaggregates might be present within the FUS condensates. SiMRoD reports on the rotational mobility of the dye probe using polarization (*p*) measurements. Millisecond-scale SiMRoD (ms-SiMRoD) is sensitive to rotational diffusion coefficient (*D_r_*) values < ∼10^3^ rad^2^/s, i.e., about four orders of magnitude slower rotational motion than measurable during the fluorescence lifetime of the fluorophore (e.g., using fluorescence anisotropy; **Fig. 5a-e**). While the effective viscosity experienced by a FUS molecule as estimated from its translational diffusion behavior will depend upon the breaking and formation of numerous interactions along the entire length of the polypeptide, the rotational motion of an probe attached via a flexible linker will be largely determined locally within the vicinity of the dye and could vary widely depending on the local solvent properties and the interactions with nearby protein neighbors. Notably, a dye attached to a translationally static molecule could still rotate relatively unhindered due to this linker, which in our case includes a two-carbon aliphatic chain between the maleimide functional group and the dye (**Fig. 1c**) as well as the lengthy disordered N-terminus of FUS. Thus, the effective viscosity for translational motion (ρι) of FUS in condensates could be largely decoupled from the effective viscosity for rotational motion of the dye (ρι_r_). At the 1 h timepoint, single molecule polarization values were narrowly distributed around 0 with a variance of Var(*p*) = 0.013 (**Fig. 5f**), indicating that *D_r_* ≥ ∼10^4^ rad^2^/s (**Fig. 5c,e**). At longer timepoints, larger *p* values were observed, indicating that some molecules had reduced rotational mobility (**Fig. 5g-j**). After 7 days of aging, the Var(*p*) = 0.027, which indicates that the average *D_r_* decreased (**Fig. 5e,j**). By comparing experimental and simulated polarization histograms, we estimate a minimum *D_r_* of ∼5×10^4^ rad^2^/s after 1 h of aging (**Extended Data Fig. 9g**). In contrast, after 7 days of aging, multiple *D_r_* values were required to fit the data, with ∼20% of dye molecules having a *D_r_* of ∼300 rad^2^/s, i.e., at least approximately two orders of magnitude slower rotational diffusion (**Extended Data Fig. 9h,i**). This implies that the ρι_r_ experienced by some of the dye molecules was > 10^5^ cP (according to the Stokes–Einstein–Debye relation for a spherical particle and a JF635b dye radius ≈ 1 nm). Recalling the flexible linker between the dye molecule and FUS, this high effective viscosity experienced by some dye molecules indicates a highly restrictive local environment, potentially indicating solid-like assemblies that could nonetheless rotationally diffuse as an ensemble. Assuming that |*p*| > 0.3 indicates a molecule with restricted rotational mobility, such immobility was identified both within cluster regions as well as outside of identified clusters (**Fig. 5k-m**). High |*p*| values outside of identified clusters do not necessarily imply the absence of a cluster, as not all clusters may have been identified or some clusters may be small. In addition, low |*p*| values (e.g., within a cluster) do not by themselves indicate high mobility, as such values are obtained for all rotational mobilities (**Fig. 5c-e**). Nonetheless, these data clearly indicate that some highly rotationally immobile dye molecules were detected within clusters. This is the expected signature of a solidified assembly.

**Fig. 5.**
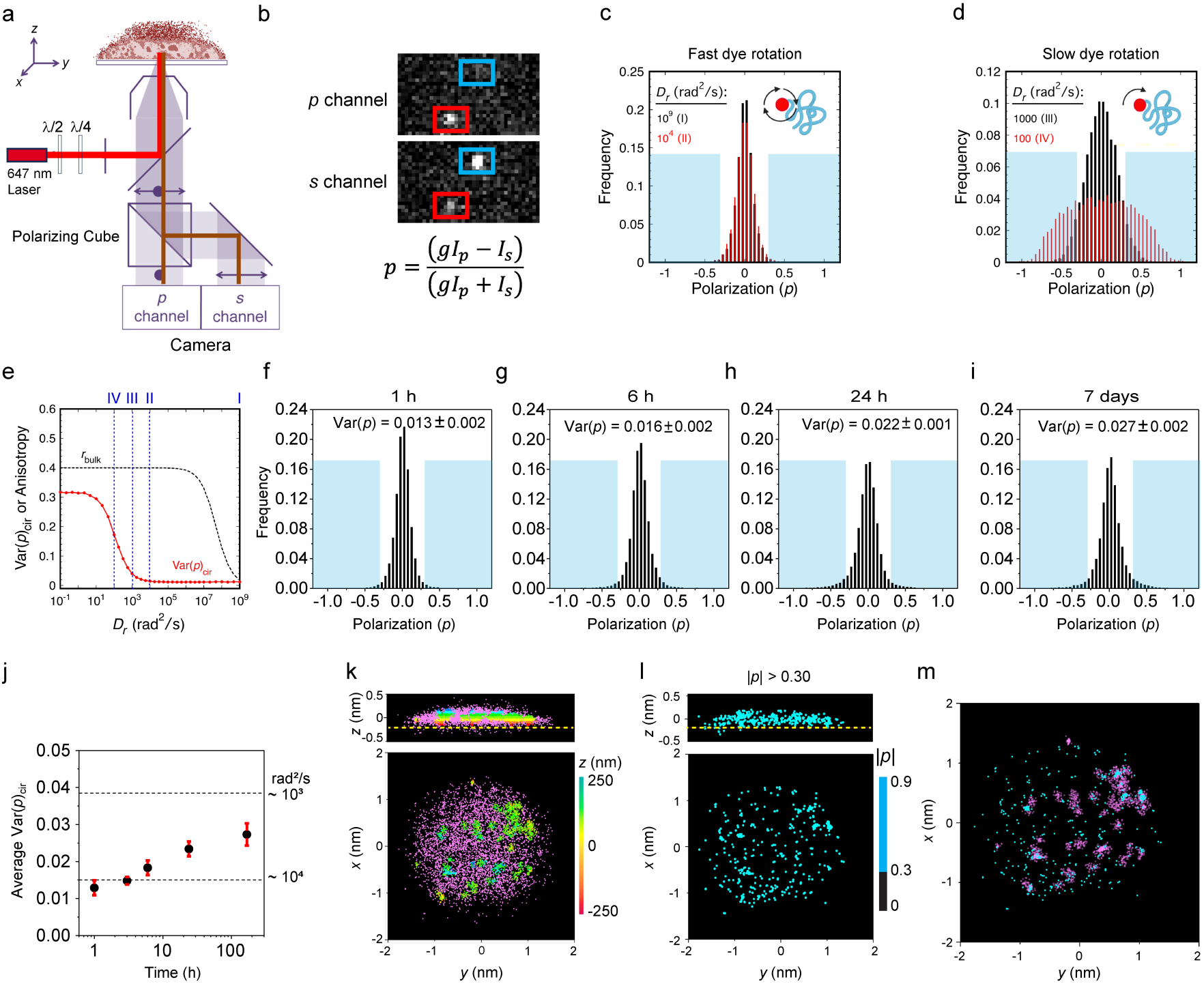
Decreased dye rotational mobility upon condensate aging. **a,b,** SiMRoD microscopy. **a,** Imaging scheme. The emission from single molecules excited by circularly polarized light is split with a polarizer and collected on two halves of a camera. **b,** Polarization calculation. The polarization is calculated from the fluorophore emission intensity measured in the two polarization channels (*I_s_* and *I_p_*), which depends on the orientation and the rotational mobility of the fluorophore. The correction factor, *g*, adjusts for the different detection efficiencies of the two channels. The spot imaged in the *blue* boxes identifies a highly immobilized fluorophore, as evidenced by the emission obtained largely in one channel. **c-e,** Simulated effects of rotational mobility on polarization measurements assuming a 5 ms imaging time. **c,d,** Polarization histograms. For rapidly rotating dye molecules, a histogram of polarization measurements is narrow, with a width determined by stochastic variations in the intensities collected and background fluctuations (**c**). For slower rotating dyes, the polarization distribution is wider due to preferred orientations during the imaging time (**d**). **e,** Effect of *D*_r_ on Var(*p*)_cir_. The width of polarization histograms using circular excitation is summarized as the variance of the polarization, Var(*p*)_cir_, and this varies depending on the rotational diffusion coefficient (*D*_r_) of the dye probe. For comparison, also shown is the bulk anisotropy (*r*_bulk_), which is sensitive to rotational mobility during the fluorescence lifetime of the probe. *n* = 10,000 for each *D*_r_ value. **f-j**, SiMRoD polarization measurements from FUS^A2C^-JF635b (1 nM) in FUS condensates at different aging times. The increased Var(*p*)_cir_ at longer aging times indicates decreased rotational mobility according to the simulations in **c-e** (5 ms/frame; *n*_1h_ =25,477, *n*_3h_ = 52,809, *n*_6h_ =17,654, *n*_24h_ = 55,747, *n*_7days_ = 28,493). See **Extended Data Fig. 9** for photon scatterplots and individual Var(*p*)_cir_ measurements. The light blue areas of the polarization histograms identify the regions of low rotational mobility, defined as |*p*| > 0.3. **k-m**, Distribution of rotational immobility within a condensate aged for 7 days. **k,** Density distribution. 3D localization information was obtained from the channel with the higher intensity (see Methods). Clusters were then identified as described in Fig. 1 and are colored based on their *z* height with all remaining localizations shown in *magenta* (*n* = 6,478 localizations). **l,** FUS^A2C^-JF635b molecules (1 nM) with |*p*| > 0.3 (*cyan*) distributed throughout the condensate in **k**. **m,** Overlap of identified clusters (*magenta*) with immobile dyes (*cyan*, |*p*| > 0.3). Approximately half (47%) of the immobile dyes were found within identified clusters.

## Discussion

This study illustrates that FUS molecules within single-component condensates exhibit multiple distinct behaviors and physical properties that vary over a distance scale of hundreds of nanometers; we denote distinctive regions of higher apparent density, lower mobility, and higher fluorescence lifetime as ‘clusters’. Such clustering intimates the potential for substantial structural and functional complexity, even for relatively simple condensates. The data reported here demonstrate the following major points. First, clusters were detectable in condensates at all timepoints, and the immobile cluster fraction increased over time. Second, immobile clusters were detected ∼0.3-0.7 µm above the coverslip surface, indicating that immobility was determined by internal structural constraints and not by adsorption to the surface. Third, at an early timepoint, the FUS molecules within clusters exhibited similar translational mobilities with variable levels of confinement, indicating diverse environmental constraints. Fourth, the average overall viscosity increased over time with a minor fraction of molecules experiencing an extremely high apparent viscosity, as inferred from rotational mobility measurements. Highly immobilized dye molecules were observed both within identified clusters and without. And fifth, while condensates never solidified, the average translational mobility decreased over time, likely a result of an increased number and/or affinity of interactions between FUS monomers. While ∼85% of FUS molecules remained mobile and migrated micrometer-scale distances in seconds, clusters became increasingly immobilized over time. One model is that a structural scaffold developed that increasingly trapped clusters over longer time period; this is consistent with the porous network structure observed earlier that permeated the condensate even at early time points^32^. Alternatively, for a viscoelastic percolated network, the effective viscosity felt by large particles can be much larger than that experienced by the monomeric components^27^. Notably, the Gibbs phase rule predicts an upper limit of two phases (e.g., dilute and condensed) at equilibrium for a single species in solvent^46^. As three distinct diffusional modes within the condensates were detected, this therefore implies a system not at equilibrium. Since the fractional composition of the diffusive species changed over time, albeit slowly, the system was semi-stable, i.e., a pseudo-equilibrium between distinct phases. While surface aggregates or a solidified shell for FUS condensates has been observed by others, we have not observed such structures^21,32,33,47^, which likely indicates a sensitivity to experimental conditions and a reflection of properties of the full-length FUS protein.

An obvious interpretation is that clusters represent distinct entities, i.e., a distinct physical state in spatially separate locales within the condensates, and the different diffusive behaviors of the labeled FUS proteins reflect partitioning into these structures. The fastest translational mobility (*D*_3_ ≈ 0.73 µm^2^/s) undoubtedly reflects the movement of the condensate structural entities themselves, i.e., FUS monomers [hydrodynamic radius in buffer between 3.7 nm^[48]^ and 3.1 nm (**Extended Data Fig. 4c**)]. Based on *D*_3_ and the trajectories themselves, a fraction of FUS explores the volume enclosed by entire condensates within seconds while experiencing an effective bulk viscosity of ∼90 cP (estimated using the Stokes-Einstein-Sutherland equation). The intermediate translational mobility (*D*_2_ ≈ 0.26 µm^2^/s) also reflects long range mobility, albeit slower, consistent with FUS monomer migration through a more connected, and hence more viscous (∼240 cP), region of the condensate. These two distinct mobility phases (*D*_3_ and *D*_2_) are both present as substantial fractions within condensates at all timepoints, including the earliest measured (1 h), although their fractions change over time, suggesting a dynamic equilibrium. The high fraction of *D*_2_ (∼69% after 7 days) suggests that this does not reflect cluster diffusion, as we do not have evidence for such a high cluster fraction. While we favor the interpretation that *D*_2_ reflects a more connected network, we cannot rule out the possibility that *D*_2_ reflects the diffusion of FUS oligomers. The low translational mobility (*D*_1_ ≈ 0.014 µm^2^/s) most certainly arises from FUS within the cluster regions. A natural assumption is that *D*_1_ simply reflects the diffusive motion of the clusters themselves (multiple hundreds of nanometers in diameter). However, such clusters with a *D*_1_ diffusion coefficient are expected to diffuse > 3 µm in a minute; this is inconsistent with the localization density maps accumulated over 20 min (**Fig. 1g,i,k**) wherein clusters were observed as static objects.

An alternate possibility is that the different particle mobilities that were observed all reflect the behavior of FUS monomers migrating through regions of different connectivities. This readily explains how single molecules can switch between distinct diffusivities and experience different levels of confinement (**Fig. 4**). In this picture, a single system-spanning percolated network does not fill the entire volume, but rather regions of different connectivities create a relatively open sponge-like structure that allows for rapid diffusion, and that also enables molecules to become immobilized or experience restricted mobility within narrower channels (**Fig. 6**). The beginnings of an interlinked scaffold structure could be assemblies of percolated molecules (‘clusters’) that can diffuse as a unit within the bulk (slow mobility within **Supplementary Videos 2-4**), and, over time, coalesce to form more elaborate structures (**Fig. 1m**). Importantly, the slow aging timescale suggests that molecules embedded within percolated structures do not need to be permanently trapped but could be stochastically released. The framework provided here is expected to guide future work in FUS and other condensates. Additional complexities are expected within multi-component condensates, such as influences on partitioning into percolated regions and how percolation is regulated.

**Fig. 6.**
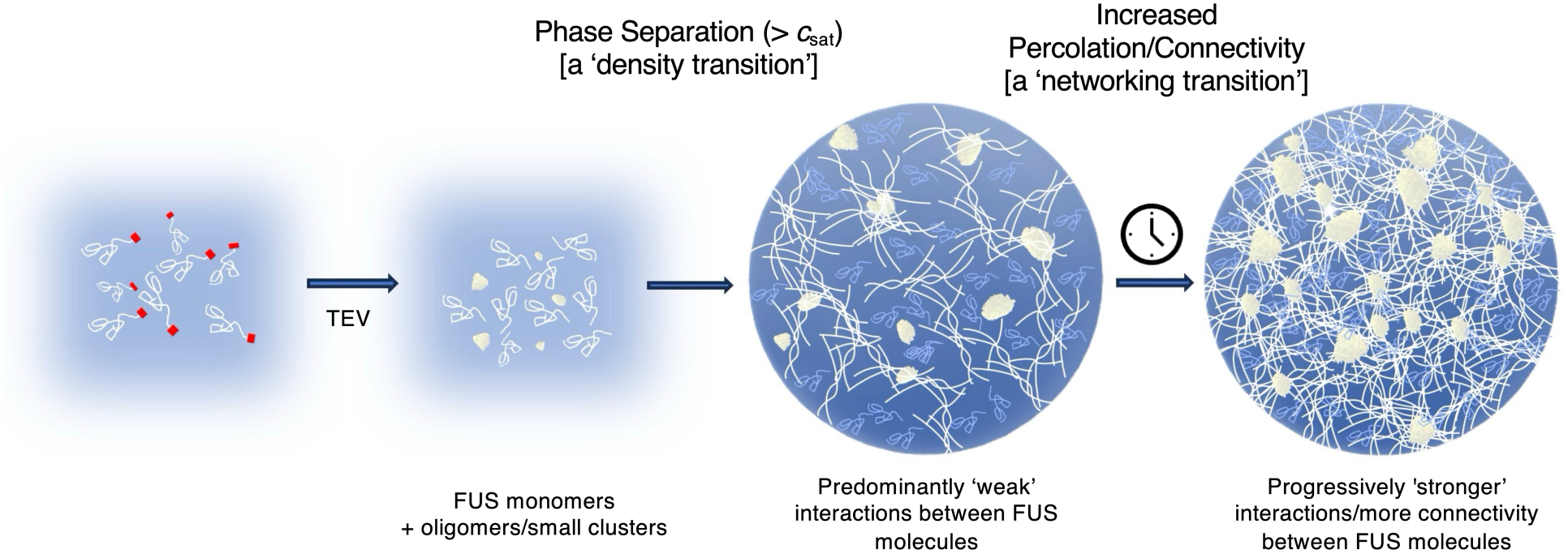
Model of Condensate Heterogeneity. Upon generation of FUS monomers by TEV protease cleavage of MBP-FUS, small oligomeric structures or clusters form, the size and distribution of which depends on physiochemical conditions. Upon reaching *c*_sat_, condensates form (a ‘density transition’) that are initially characterized by predominantly weak interactions, though even at early time points, intra-condensate clusters form that are characterized by high connectivity. These clusters are initially predominantly mobile, though an internal network structure can immobilize some of them. Over time, an increase in overall connectivity, potentially due to conformational rearrangements, increases network interconnectivity (a ‘networking transition’) and leads to increased immobility of the clusters while allowing for high mobility of the majority of the FUS molecules.

## Methods

### Overproduction and purification of FUS variants

The MBP-FUS and MBP-FUS^A2C^ proteins contain an N-terminal maltose binding protein (MBP) tag for retaining solubility and a C-terminal 6xHis-tag for purification (**Fig. 1a**). Both the MBP and 6xHis-tags are removed by TEV protease activity. MBP-FUS and MBP-FUS^A2C^ were overproduced from plasmids pMBP-*tev*-FUS-*tev*-His_6_ (Addgene #242384) and pMBP-*tev*-FUS^A2C^-*tev*-His_6_ (Addgene #242385), respectively. These plasmids were generated by PCR amplification of the FUS coding sequence in pMBP-*tev*-FUS-EGFP-*tev*-His_6_ [*aka* pMal-*Tev*-FUS(WT)-EGFP-*Tev*-His_6_, as described in Hofweber et al.^49^] with SalI/HindIII ends without or with the A2C mutation and ligating the digested fragment into the original plasmid digested with SalI/HindIII, which removes the FUS-EGFP coding sequence. Coding sequences were verified by DNA sequencing. Plasmid constructions are described in the history of the SnapGene files available on Addgene. MBP-FUS and MBP-FUS^A2C^ were overproduced and purified essentially as described^34^, with minor modifications. For completeness, the complete procedure is provided below.

FUS variants were overproduced in *E. coli* strain Rosetta 2(DE3)pLysS (Novagen) transformed with the appropriate plasmid using lysogeny broth (LB) with 100 µg/mL ampicillin and 34 µg/mL chloramphenicol. After incubating a 20 mL starter culture overnight at 37 °C, it was centrifuged (4000×*g*, 5 min, 25 °C) and resuspended in 4 mL fresh media. Resuspended culture (1 mL each) was added to 4×500 mL fresh LB + antibiotics. Cultures were incubated at 37 °C until an OD_600_ of ∼0.8. Protein overproduction was induced with 1 mM isopropyl β-D-1-thiogalactopyranoside (IPTG) for 24 h at 12 °C with shaking at 250 rpm. Cells were recovered by centrifugation (4,000×*g*, 20 min, 4 °C) and resuspended in 40 mL resuspension buffer [50 mM Na_2_HPO_4_/NaH_2_PO_4_, 1 M NaCl, 10 μM ZnCl_2_, 10 mM imidazole, 4 mM β-mercaptoethanol (βME), 10% glycerol, pH 8.0], and protease inhibitors [10 mM phenylmethane sulfonyl fluoride (PMSF), 100 µg/mL trypsin inhibitor, 20 µg/mL leupeptin, and 100 µg/mL pepstatin]. The suspension was lysed by French press (3× at 16,000 psi), the lysate was centrifuged (50,000×*g*, 20 min, 4 °C), and the supernatant was incubated with 4 mL of Ni-NTA resin (Qiagen), that was prewashed and equilibrated with 10 column volumes (CVs) of lysis buffer, for 30 min at 25 °C. The suspension was transferred to a gravity column. The resin was washed with 10 CVs of wash buffer A (50 mM Na_2_HPO_4_/NaH_2_PO_4_, 500 mM NaCl, 10 μM ZnCl_2_, 10 mM imidazole, 4 mM βME, 10% glycerol, pH 8.0). Fractions (1 mL) were eluted with elution buffer A (50 mM Na_2_HPO_4_/NaH_2_PO_4_, 500 mM NaCl, 10 μM ZnCl_2_, 300 mM imidazole, 4 mM βME, 10% glycerol, pH 8.0). Protein-containing fractions (∼15 mL) were combined and diluted to 100 mL using dilution buffer (50 mM Na_2_HPO_4_/NaH_2_PO_4_, 150 mM NaCl, 10 μM ZnCl_2_, 10 mM imidazole, 10% glycerol, 4 mM βME, pH 8.0). After gentle shaking at room temperature for 5 min, the protein mixture was loaded onto a gravity flow column with 2 mL amylose resin (New England Biolabs, #E8021) that was prewashed and equilibrated with 10 CVs of dilution buffer. The resin was washed with 5 CVs of wash buffer B (50 mM Na_2_HPO_4_/NaH_2_PO_4_, 200 mM NaCl, 10 μM ZnCl_2_, 10 mM imidazole, 5% glycerol, pH 8.0), and the protein was eluted with amylose elution buffer (20 mM Tris, 150 mM NaCl, 20 mM maltose, 10 μM ZnCl_2_, 5% glycerol, pH 8.0). For MBP-FUS^A2C^, 4 mM tris(2-carboxyethyl)phosphine (TCEP) was used in place of βME in wash buffer B and 1 mM TCEP was included in the amylose elution buffer. Purified proteins were flash frozen in liquid nitrogen immediately after purification and stored at −80°C in 25 µL aliquots until use.

### Dye labeling of MBP-FUS^A2C^

MBP-FUS^A2C^ was incubated with a 15- to 20-fold molar excess of JF635b-maleimide (Bio-techne|Tocris, #8156), JF549-maleimide (Bio-techne|Tocris, #6500), or CAGE635-maleimide (Abberior, #CA635-0003) at room temperature for 15-20 min. The reaction was quenched with 1 mM βME. The dye-protein mixture was incubated for ∼30 min with 0.1 mL Ni-NTA resin [prewashed and equilibrated with 10 CVs of equilibration buffer (20 mM Tris, 150 mM NaCl, 10 μM ZnCl_2_, 5% glycerol, pH 8.0)] and then loaded into a gravity flow column. Excess dye was removed by washing the resin-bound protein with 100 mL of wash buffer C (20 mM Tris, 1 M NaCl, 0.1% Triton X-100, 5% glycerol, pH 8.0) and 50 mL of wash buffer D (20 mM Tris, 150 M NaCl, 10 mM imidazole, 5% glycerol, pH 8.0). The protein was eluted (250 µL fractions) with elution buffer B (20 mM Tris, 300 mM imidazole, 150 mM NaCl, 5% glycerol, pH 8.0). Dye-labeled proteins were flash frozen in liquid nitrogen and stored at −80 °C in 3 µL aliquots until use.

### Protein concentration and purity

Protein concentrations (typically ∼15–30 µM for MBP-FUS and 1-2 µM for dye-labeled MBP-FUS^A2C^) were determined by the densitometry of bands on SDS-PAGE gels stained with Coomassie Blue R-250 using carbonic anhydrase as a standard and a ChemiDoc MP imaging system (Bio-Rad Laboratories). The purity of dye-labeled proteins was assayed by direct in-gel fluorescence imaging using the same ChemiDoc MP imaging system and was determined to be > 95%. The JF635b dye was visualized after soaking the acrylamide gel in ∼5% acetic acid for 5 min to convert the blinking dye molecules into the protonated fluorescent state. The CAGE635 dye was visualized after a brief illumination with UV light to photoactivate the dye molecules.

### FUS condensate formation and sample preparation

FUS protein stock solutions were thawed at room temperature and centrifuged (14,000×*g*, 10 min, 25 °C). Supernatants were used immediately. MBP-FUS (14 µM final) and dye-labeled MBP-FUS^A2C^ (1-20 nM final) were mixed in droplet buffer A (20 mM Na_2_HPO_4_/NaH_2_PO_4_, 2.5% glycerol, 150 mM NaCl, pH 7.5). Protein mixtures (5 µL) were then diluted 2-fold with 4.7 µL of no salt droplet buffer B (20 mM Na_2_HPO_4_/NaH_2_PO_4_, 2.5% glycerol, pH 7.5) and 0.3 µL (3 U) of AcTEV protease (10 U/µL; Invitrogen, #12575015) to yield a final NaCl concentration of 75 mM and 7 µM MBP-FUS. AcTEV protease activity results in removal of the MBP solubility tag and the 6xHis-tag^49^, thus initiating condensate formation (**Extended Data Fig. 1**). After incubation at room temperature for 10 min, the solution containing condensates was added to a microscope slide flow chamber (∼10 µL), which was then sealed with clear nail polish or Eco-sil Soft silicone (Picodent #13008100) to avoid evaporation. Flow chambers were made as follows. Large coverslips (Gold Seal 3243; 24×60 mm; ThermoFisher Scientific, #50-189-9138) were plasma cleaned. Small coverslips (10.5×22 mm; Electron Microscopy Sciences, #72191-22) with double-sided tape on their short edges were pasted on top of a large coverslip to construct flow chambers. The flow chambers were then treated with 2% mPEG-silane (LaysanBio, #MPEG-SIL-5000) for ∼45 min at room temperature to passivate the surface, followed by 3×10 µL washes with droplet buffer C (20 mM Na_2_HPO_4_/NaH_2_PO_4_, 2.5% glycerol, 75 mM NaCl, pH 7.5).

### Narrow-field imaging

Condensates containing 0.5 nM FUS^A2C^-JF549 were added to flow chambers for imaging. A Nikon Eclipse Ti microscope equipped with a 100X oil-immersion objective (Nikon Apo TIRF, 1.49 NA) was used to image via narrow-field epifluorescence excitation^50^ with 561 nm light. Images were magnified (1.5X tube lens) and recorded using a CMOS camera (Prime 95B; Photometrics) at 200 ms/frame. Images were analyzed for colocalization using the coloc2 plugin in Fiji^51^.

### FLIM and FCS

Fluorescence Lifetime Imaging Microscopy (FLIM) and Fluorescence Correlation Spectroscopy (FCS) experiments were performed on a Luminosa time-correlated single photon counting confocal microscope system (Picoquant). For FLIM, condensates containing 10 nM FUS^A2C^-JF549 or 10 nM of FUS^A2C^-CAGE635 were imaged by zooming in on the condensates and acquiring 128×128 images for ∼3 min to capture sufficient photons for a reliable multi-component lifetime fit (until 4000 photons acquired in the brightest pixel). Average weighted lifetimes were determined by combining all the arrival time data from a condensate and fitting with a double exponential fit using SymPhoTime (Picoquant). FLIM images show the amplitude-weighted average fluorescence lifetime per pixel. FCS measurements were performed on purified proteins using the system software to fit FCS curves.

### Dynamic light scattering (DLS)

DLS experiments on FUS variants were performed on a Zetasizer Nano (Malvern Panalytical) using a minimum concentration and volume of 0.5 µM and 100 µL, respectively.

### 3D super-resolution imaging using astigmatism microscopy

**Microscope system.** The set-up, sensitivity, and precision of our microscope system using a tunable astigmatism to localize single fluorophores in three-dimensions (3D) was described previously^35,36,52^. Briefly, a Zeiss Axiovert 200M equipped with an oil-immersion objective (Zeiss Alpha Plan-Apochromat, 1.40 NA) and Kinetix sCMOS camera (Teledyne Photometrics; 2×2 pixel binning yielded square pixel dimension = 138 nm at the image plane) were used to capture the fluorescence emission from individual JF635b or CAGE635 molecules during their on-time events within FUS condensates. Emission intensities were converted to photons using an analog to digital conversion (ADC) factor of 0.27 during image analysis. Laser excitation was converted to circularly polarized light with half (λ/2) and a quarter (λ/4) waveplates (Newport, #10RP42-1 and #10RP44-1) before entering the microscope objective as a slightly converging illumination beam. A self-configured adaptive optics system (Imagine Optic, AOKit Bio) incorporated into the emission path generated the *z*-dependent spot ellipticity needed for 3D information using 60 nm root mean square (rms) astigmatism^36^. A self-configured TIRF-Lock system (Mad City Labs) provided *z*-stability for the coverslip of < 3 nm for the duration of the experiment.

Image sequences were analyzed using the ThunderSTORM plug-in^53^ for Fiji^51^. The maximum likelihood method was used to determine all single molecule spot centroids (*x*, *y*) and the “elliptical gaussian” fitting option was used to fit the astigmatic point spread functions (PSFs), which yielded the *z*-positions. The calibration file required for the *z*-dependent PSF was obtained from *z*-stacks (−500 nm to +500 nm; 25 nm steps) of 0.1 µm fluorescent microspheres (ThermoFisher Scientific, T7279). These data were fit as described by Huang *et al.*^54^ and implemented in the ThunderSTORM plug-in. The post-processing option “remove duplicates” was used to eliminate molecules repeated in successive frames.

**Simulations.** Simulated random distributions of points and jump steps were generated by **Jump Step Histogram Simulator2**, a lab-constructed Microsoft Excel spreadsheet that simulates jump step histograms for a population of particles with up to three diffusion coefficients and with astigmatism-based localization precisions. For these simulations, the *z*-dependent localizations precisions (α*_x_*, α*_y_*, and α*_z_*) were defined by experimental calibration curves^36^ using a randomly determined photon count based on experimentally-determined photon distribution histograms (see **Extended Data Fig. 2a,b**). The initial point was chosen randomly from within a user-defined rectangular box or cylinder. The final point was determined after one translational step each in *x*, *y*, and *z* selected randomly from a normal distribution centered around the starting position with a step variance (Var) of 2*Dt* in each direction, where *D* is the diffusion coefficient and *t* is the time for each step (imaging time). Measurement error (localization precision) was added to each localization randomly selected from a normal distribution with a standard deviation equal to the localization precision, and then, the distance (*R*) between the simulated measurement values was calculated. A histogram of such *R* values is a simulated jump step histogram (e.g., **Fig. 3b**). The distributions in **Fig. 1h,j,l** were obtained from the initial points in such simulations. The program calculates the average precisions under the input conditions; this feature was used to estimate the average localization precisions reported for astigmatism-based imaging in this paper.

**Density maps.** To generate 3D density maps, condensates were formed with 5 or 10 nM FUS^A2C^-JF635b. For the 24 h and 7 day timepoints, the JF635b blinking frequency was slightly higher leading to increased background. Therefore, 5 nM FUS^A2C^-JF635b was used for the longer timepoints. Localizations (one localization/blink) were acquired with 647 nm illumination using ∼10 kW/cm^2^ at the sample and 30 ms/frame for 20 min. Only localizations with > 500 photons and |*z*| < 500 nm were retained. The resultant average localization precisions for the experiments were σ_x_ = 13.4 nm, σ_y_ = 15.6 nm and σ_z_ = 22.8 nm, which were determined as described earlier^35^ by **Jump Step Histogram Simulator2** (see **Extended Data Fig. 2a,b**).

To identify regions of higher localization densities within condensates, the DBSCAN algorithm^55^ was implemented in MATLAB. The distance threshold ‘ε’ required for DBSCAN was calculated from using MATLAB’s *k*-nearest neighbors (kNN) algorithm on simulated random distributions (from **Jump Step Histogram Simulator2**) at equivalent densities over the entire volume. The kNN algorithm estimates the distance to the k^th^-nearest neighbor and ε was estimated based on the knee of the kNN output curve, as calculated by MATLAB (see **Extended Data Fig. 2f**). The “minimum points” input for DBSCAN was chosen as the value that yielded no clusters with the respective ε from the corresponding simulated random distribution. To estimate the size of clusters identified by DBSCAN, individual clusters were isolated after visualizing them in 3D. The isolated clusters were fit to an ellipsoid to estimate the lengths of the three principal axes, which were averaged to give the average principal axis length.

To simulate localization distributions at the same average density of the experimental measurements, two methods were compared to estimate the volume of the condensate occupied by the measured localizations. For the first method (‘cube volume estimator’), the observation volume was subdivided into cubes with sides of 100 nm or 125 nm, and then the number of cubes with at least one localization was determined. The number of non-empty cubes times the cube volume yielded the estimated volume of the condensate. The number of localizations obtained from the condensate was divided by the estimated volume to obtain the estimated localization density (in localizations/µm^3^) for the condensate, which provides a normalization for easier comparison of condensates of different volumes. Since the number of localizations obtained from a condensate is a fixed number, the estimated volume of the condensate directly influences the estimated localization density. Hence, to estimate the error of the cube volume estimator method, random localizations within a cylinder of varying density were simulated and the resultant ‘estimated density’ was calculated. For comparison, MATLAB’s convex_hull algorithm (convhull), which estimates the size of irregular shaped objects, was also used to estimate the occupied volume, which was converted to estimated density for comparison (see **Extended Data Fig. 2d**).

**3D tracking.** FUS condensates with 1 nM FUS^A2C^-CAGE635 were imaged at 5 ms/frame for ∼25 min with intermittent bursts of UV illumination (405 nm) to activate the CAGE635 fluorophores (∼1.5 kW/cm^2^ for 5 s every 10,000 frames). Spots were localized using ThunderSTORM as described earlier. Only localizations with > 500 photons (see **Extended Data Fig. 6a**), and |*z*| < 250 nm were included in the analysis. The average precisions were σ_x_ = 12.7 nm, σ_y_ = 15 nm and σ_z_ = 21 nm, which were determined as described earlier^35^ by **Jump Step Histogram Simulator2**. To perform tracking with high confidence and eliminate the potential for errors arising from linking successive localizations into tracks, we removed frames that contained multiple spots. Only tracks with ≥ 5 localizations were used to build jump step histograms.

**Jump step histogram analysis.** To identify confined trajectories within condensates from astigmatism data (e.g., **Fig. 3a**), we eliminated those tracks that had an average jump step > 50 nm. All remaining tracks were confirmed visually to exhibit confined motion. Jump histograms from these tracks (e.g., **Fig. 3b**) were fit to^35^:

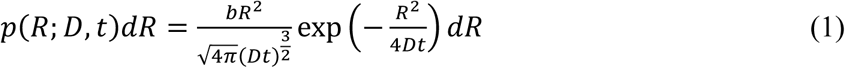

where *t* is the time between successive localizations, *b* is the bin size of the jump distances, *R* is the three-dimensional jump step, and *D* is the diffusion coefficient. However, to obtain an accurate diffusion coefficient, the influence of localization precision needs to be accounted for. To achieve this, jump step histograms were simulated using a MATLAB version of **Jump Step Histogram Simulator2** named **Jump Step Histogram Fitter**, which enabled determination of the best fit from the lowest root mean squared deviation (RMSD) via a built-in genetic algorithm.

To fit jump histograms obtained from all tracks, a jump step histogram including three diffusion coefficients was needed^35^:

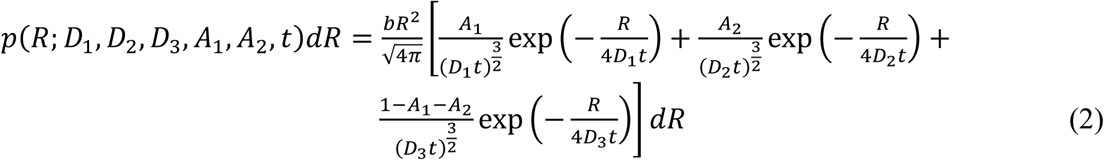

where *A*_1_ and *A*_2_ are weighting factors for the three distributions arising from the diffusion coefficients *D*_1_, *D*_2_, and *D*_3_. The fractions of molecules with these three diffusion coefficients are *A*_1_, *A*_2_, and *A*_3_ (= 1 - *A*_1_ - *A*_2_), respectively (see **Fig. 3e**). Here too, the influence of localization precision needs to be accounted for, and so a direct fit using **Eq. 2** would over-estimate the *D* values. Recognizing that at least a two *D* fit was required (the confined molecules clearly exhibited smaller jump steps than the majority of molecules – see **Fig. 3**), we tested both two and three *D* models. The three *D* model worked better (see **Extended Data Fig. 6d-f**), and was therefore used to estimate the parameters for all timepoints (**Fig. 3d,e**). **Jump Step Histogram Fitter** was implemented assuming a range of 0-0.02 µm^2^/s for *D*_1_ (based on **Fig. 3b**), 0-0.5 µm^2^/s for *D*_2_, and 0.5-1 µm^2^/s for *D*_3_.

### SiMRoD microscopy

FUS condensates can mature over time from a high mobility liquid state to a low mobility solid state^38,56^. This phase maturation can, in principle, be detected by the change in rotational mobility of a fluorescent dye attached to FUS molecules. In our case, the JF635b dye was attached via a flexible linker to an intrinsically disordered region of FUS^57^. Thus, rotational mobility measurements were expected to depend predominantly on the local environment of the dye instead of the rotational motion of the macromolecule to which it was attached, although these mobilities could be interlinked. Probes within a completely solid structure attached to the surface of the chamber are expected to be rotationally immobile on any timescale. In contrast, probes within a large solid assembly (i.e., an aggregated particle) diffusing in a viscous but liquid-like medium would likely be rotationally immobile in a fluorescence anisotropy experiment but have rotational mobilities distinct from a completely solid structure. Single molecule rotational diffusion (SiMRoD) microscopy, originally reported as polarization PALM or ‘p-PALM’^45^ and renamed here, can distinguish such slow rotational mobilities. Since SiMRoD is a relatively new technique, we first describe the basic principles of the approach before describing the measurements on FUS condensates.

In camera-based SiMRoD, the fluorescence emission from a single fluorescent molecule is split with a 50% polarizing beam splitter and both emission channels are simultaneously recorded^45^. Single molecule polarization values, *p*, are calculated from the spot intensity pairs using:

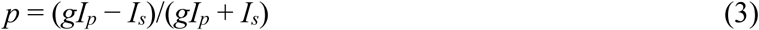

where *I_p_* and *I_s_* are the intensities (photons) measured in the two detection channels, and *g* is a system-dependent factor that corrects for differences in photon collection efficiency of the *p*- and *s*-channels. Each individual polarization value estimates the time average of the polarizations from the rotational trajectory of the individual molecule during the acquisition time. Thus, *p* values near zero are expected for fast rotational motion relative to the image collection timescale. In contrast, *p* values ranging from −1 to 1 are expected for immobile molecules. In SiMRoD, average rotational mobility information is extracted not from the average bulk polarization, but from polarization measurements obtained from thousands of individual molecules that are pooled to generate polarization frequency histograms (**Fig. 5c,d**). The primary experimental readouts from these data are the average polarization, <*p*>, and the variance of the polarization distribution, Var(*p*). The overall shape of polarization histograms and photon scatterplots can provide additional clues as to the underlying physical constraints on the probe’s rotational mobility^45^. Var(*p*) provides a measure of the width of polarization histograms (see **Fig. 5e**), and is formally calculated using:

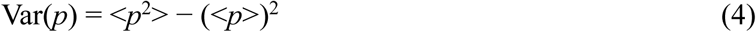

where <*p*^2^> is the average square of the measured polarization values. For a properly calibrated system using circularly polarized excitation,

<*p*> = 0 and hence Var(*p*) = <*p*^2^>. Thus, we assumed that Var(*p*)_cir_ was equivalent to the average of the squares of the experimentally determined *p* values, <*p*^2^>_cir_. The requirement that <*p*>_cir_ = 0 (except under unusual circumstances) provides a convenient internal check for each dataset on instrument alignment and calibration.

For SiMRoD experiments in condensates containing FUS^A2C^-JF635b, the emission from a single emitter was split with a 50% polarizing beam splitter cube (PBS201, ThorLabs, Newton, NJ) mounted in an Optosplit III beamsplitter (Cairn Research, Kent, UK), which allowed the two polarization components to be imaged simultaneously on the two halves of the Kinetix sCMOS camera. For circular excitation, <*p*> is expected to be 0. Therefore, according to **Eq. 3**, *g* = <*I_s_*>/<*I_p_*>, which was experimentally measured for each condensate independently by overlapping the photon distribution curves from both channels. The value of *g* was considered optimal when the photon distribution curves from the *p*- and *s*-channels overlapped with each other. Across all timepoints of condensate measurements, *g* ranged from 0.90-1.00. Since SiMRoD depends only on the intensity (photons) measured in the two channels, it is insensitive to the astigmatic PSFs or minor spatial misalignments of the two channels. Data were collected at 5 ms/frame for 8 min using ∼50 kW/cm^2^ (λ_ex_ = 647 nm), measured at the sample plane.

### Simultaneous SiMRoD and 3D localization

For 3D SiMRoD, the two images needed for SiMRoD polarization measurements both contained astigmatic spots obtained with the Optosplit beamsplitter on the astigmatism microscope described earlier. 3D localization information was obtained from the higher intensity channel, with all such *s*-channel localizations translated to the reference *p*-channel. The *p*- and *s*-channels were first spatially aligned by identifying all emitters that yielded at least 500 photons in both polarization channels and then calculating the translation matrix (using the MATLAB program Displacement.m) that yielded the minimum average displacement of the centroids of the emission patterns obtained in both channels. Using this alignment matrix, all localizations within spot pairs that had higher-intensities in the *s*-channel were then translated to the *p*-channel. The 3D coordinates were then determined for each spot pair using the intensity pattern from the channel with the higher total photons. Polarization values were then determined by **Eq. 3**.

### SiMRoD Simulations

SiMRoD simulations were performed as described earlier^45^ assuming an image integration time of 5 ms, a numerical aperture of 1.46, a photon threshold in the *p*- or *s*-channel of 200 photons, a fluorescence lifetime of 3.5 ns (which has no effect on <*p*>_cir_), a noise level of 63 photons per 12×12 pixel region of interest, and 4200 rotational walk steps (yielding an average of ∼1150-1300 photons/measurement).

### 3D tracking using MINFLUX

**Microscope system.** The MINFLUX instrument used has been described in detail^52^. In short, a 100x oil immersion objective lens (UPL SAPO100XO/1.4, Olympus, Tokyo, Japan) and 642 nm CW excitation laser was used for FUS^A2C^-CAGE635 tracking. Two avalanche photodiodes (SPCM-AQRH-13, Excelitas Technologies, Mississauga, Canada) with detection ranges of 650-685 nm (red channel) and 685-760 nm (far red channel) were used with a pinhole size corresponding to 0.78 airy units. All hardware was controlled by Abberior Imspector software (version 16.3.13924-m2112). Drift during tracking was minimized using the built-in stabilization system with typical drifts < 1 nm in *xyz*. Scattering from 200 nm gold nanoparticles (Nanopartz, Cat# A11-200-CIT-DIH-1-10) pre-deposited on the coverslip surface were used as a positional reference for the sample stabilization. A *z*-scaling factor of 0.67 was used^52^.

**Imaging.** Slides and coverslips used for MINFLUX were cleaned using a plasma cleaner, and flow chambers were made using double-sided sticky tape, as for astigmatism imaging. Undiluted 200 nm gold nanoparticles (Nanopartz, #A11-200-CIT-DIH-1-10) were added to each channel, and 15 min later the unbound particles were washed away with 1X PBS. Condensates with 1 nM FUS^A2C^-CAGE635 were then added to the flow chambers and sealed with Eco-sil Soft silicone. Condensates that were aged for ∼3-4 h were imaged for ∼2-2.5 hours, yielding an average of ∼800 tracks per condensate. Non-default measurement parameters for the individual MINFLUX scan iterations are summarized in **Supplementary Table 1**. The CFR (center frequency ratio), DCR (detector channel ratio), and EFO (effective frequency at offset) collected during MINFLUX acquisition were used for data filtering. These parameters and how they are used were described previously^52^. Briefly, a high CFR suggests the presence of a second fluorophore, and the EFO can be used to distinguish background from one or two fluorophores of interest (see **Extended Data Fig. 7a-c**). Both the CFR and EFO were obtained by combining the photons collected in the red (650-685 nm) and far-red (685-760 nm) channels. The DCR measures the photons collected in the red channel divided by the photons collected in the far-red channel. Tracks were chosen only if they had an average CFR < 0.8, a total of ≥ 90% of CFR values < 0.8, and an EFO between 150 kHz and 250 kHz. Individual localizations at the extremities of the track were discarded if CFR > 0.8. Multiple successive CFR values > 0.8, with at least one value > 1.0 was sufficient to end a track or split it into two separate tracks. Jump step histograms from filtered tracks were fit by comparison with simulated data (described in the next paragraph). Mean squared displacement (MSD) curves were calculated only from tracks that had ≥ 150 localizations. Tracks with similar MSD curves were averaged, and only the first third of these averaged trajectories are shown as this is the part of the MSD curve deemed reliable^58^ (**Fig. 4d**). To quantify track confinement, the averaged MSD curves were fit to the following^59^ to estimate α, which is a measure of the anomalous nature of the diffusion:

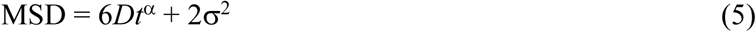

The second term is a measure of the localization error^60^, where α^2^ = α_x_^2^ + α_y_^2^ + α_z_^2^. This formula ignores the effect of motion blur, which has not been determined for MINFLUX imaging, and results in an underestimate of *D* (by < 20%) for freely diffusing molecules^61^. The confined molecules have minimal motion blur. Assuming that α_x_ ≈ α_y_ = 1.6α_z_ based on earlier results^52^, then α_x_ = α/1.54.

**Simulations.** Simulated trajectories were generated by **Trajectory Simulator**, a lab-constructed Microsoft Excel spreadsheet that generates particle trajectories exhibiting up to three diffusion coefficients and user-defined constant localization precisions in *x*, *y*, and *z*. The initial point was chosen as the center of or a random location within a user-defined sphere. Subsequent points were determined after a translational step each in *x*, *y*, and *z* selected randomly from a normal distribution centered around the starting position with a step variance (Var) of 2*Dt* in each direction. The simulation allows for molecules with three distinct diffusion coefficients (only two were used) and the molecule could be allowed to switch between diffusion modes within a trajectory or not. To prevent escape from a small spherical confinement zone, translational steps were divided into 10 sub-steps (user-definable), and the molecule could be allowed to switch between diffusion modes for each sub-step or not. If a step would allow a molecule to escape from the defined spherical volume, the step was retried up to 9 times. If all ten trials failed, the trajectory was rejected. Measurement error (localization precision) was added to each localization randomly selected from a normal distribution with a standard deviation equal to the user-defined localization precisions. The distances (*R*) between measurements were calculated to generate a jump step histogram (see **Extended Data Fig. 7f,g**). Simulated trajectories were compared with MINFLUX tracks (**Fig. 4b**).

## Data Availability

All data described in the article are shown in the figures and provided in the Supplementary Information. Source data for the main figures and Extended Data figures are provided as Supplementary Information.

## Code Availability

Data were curated and analyzed with custom code written in MATLAB R2021b. This custom code was essential to the analysis and central for extracting conclusions from the data. All MATLAB scripts used for data analysis, along with default parameters and model data, are available in the GitHub repository (https://github.com/npctat2021/3DImaging_FUS-Condensates). A user guide for the scripts is provided in the repository’s ‘README’ section. FCS and FLIM data were acquired using the Luminosa system software (v1.5) and analyzed with SymPhoTime 64 (v2.7), both provided by PicoQuant GmbH, Germany. The following commercial software packages were used for data analysis, data presentation and simulations: Imspector 16.3.15620-m2205-MINFLUX_BASE, ParaView 5.8.1, Image J (Fiji 1.52P), Origin 8.5, Kaleidagraph 5.01 and Microsoft Excel 16.76 (23081101). The Microsoft Excel spreadsheet used for simulating jump steps (Jump Step Histogram Simulator2.xlsx) is provided in figshare.

## Acknowledgements

Thanks to E. Lemke (Mainz, Germany) and J. Mittal (Texas A&M Univ) for critical comments. JF635b-maleimide was a gift from L. Lavis (Janelia, VA) prior to publication and commercial release. Plasmid pRSET-mEosEM was a gift from P. Xu (Addgene ID#132708). This research was supported by grants from the NIH (GM126190 and NS141768 to SMM), The Welch Foundation (BE-1962 to SMM), a SEED Grant from GITAM (2024/0366 to RC), and a Prime Minister Early Career Research Grant from the Anusandhan National Research Foundation (ANRF/ECRG/2024/002373/CS to RC). The authors acknowledge the assistance of the Joint Microscopy Laboratory in the Texas A&M University College of Medicine and the NIH for instrument funding support of a Luminosa confocal microscope (S10 OD032208). The authors acknowledge the access and services provided by the Imaging Centre at the European Molecular Biology Laboratory (EMBL IC), generously supported by the Boehringer Ingelheim Foundation.

## Author contributions

S.M.M., A. Sharma, and A. Sau conceptualized the project, analyzed the data and interpreted the experimental results, with contributions from other authors. A. Sharma and S.S. conducted the MINFLUX data acquisitions. A. Sharma and R. Chowdhury developed and automated the MATLAB analysis scripts. S.D. developed plasmid constructs, protein purification, dye labeling, slide preparation and condensate forming protocols, with assistance from S.H. and D.D. Initial data acquisition protocols were developed by S.D. and A. Sau. The FCS and FLIM experiments were performed by S.R. and A. Sharma. The manuscript was drafted by A. Sharma, A. Sau and S.M.M., with contributions from all other authors. All authors contributed to the revision process and approved the final manuscript. S.M.M. supervised and administered the project and garnered the primary funding to support the work.

## Additional information

**Supplementary information** The online version contains supplementary material available at https://????.

**Correspondence and requests for materials** should be addressed to Siegfried Musser.

## Peer review information

**Reprints and permissions** information is available at http://????.

---

**Supplementary Table 1.** Parameters used for FUS^A2C^-CAGE635 3D MINFLUX tracking.

|  | Iteration* |  |  |  |  |
| --- | --- | --- | --- | --- | --- |
|  | 1<sup>st</sup> | 2<sup>nd</sup> | 3<sup>rd</sup> | 4<sup>th</sup> | 5<sup>th</sup> |
| ***L* size (nm)** | 288 | 1440 | 288 | 151.2 | 100.8 |
| **TCP** | Hexagon | Zline | Octahedron | Octahedron | Octahedron |
| **Minimum number of collected photons** | 20 | 300 | 40 | 30 | 30 |
| **647 nm laser power\*\*** | 1x | 1x | 1x | 2.5x | 4.5x |
| **Minimum TCP dwell time (ms)** | 0.5 | 2 | 0.2 | 0.2 | 1 |
| **Pattern repeat** | 1 | 1 | 1 | 1 | 1 |
| **CFR check** |  |  | 0.5 |  | 3 |
| **Background threshold (kHz)** | 30 | 30 | 10 | 35 | 30 |

\*Iterations 1-4 were used for initial fluorescent particle localization and the 5<sup>th</sup> iteration was used to continuously locate the particle until it was lost.
\*\*1x = 128.3 μW, measured at the sample plane.

**Supplementary Table 2.** Volume and density values of all condensates used to construct droplet density maps in Fig. 1g-l and Extended Data Fig. 3.

| Time | Figure | Volume ($\mu\text{m}^3$) | Total Localizations | Average Density (Localizations/$\mu\text{m}^3$) | Percent of Localizations within Clusters |
| --- | --- | --- | --- | --- | --- |
| **1 h** | Fig. 1g | 7.06 | 12,717 | 1,800 | 1.6 |
|  | ED Fig. 2a (left) | 4.08 | 8,820 | 2,162 | 1.8 |
|  | ED Fig. 2a (right) | 2.43 | 5,236 | 2,152 | 2.5 |
| **6 h** | ED Fig. 2b (left) | 4.73 | 13,698 | 2,899 | 3.2 |
|  | ED Fig. 2b (right) | 10.70 | 22,203 | 2,168 | 4.4 |
|  | Image Not Shown | 2.59 | 6,483 | 2,500 | 3.3 |
| **24 h** | Fig. 1i | 5.21 | 17,179 | 3,294 | 5.7 |
|  | ED Fig. 2c (left) | 18.15 | 54,050 | 2,976 | 4.8 |
|  | ED Fig. 2c (right) | 4.26 | 17,906 | 4,200 | 6.3 |
| **7 days** | Fig. 1k | 8.20 | 25,993 | 3,169 | 21.4 |
|  | ED Fig. 2d (left) | 7.20 | 19,450 | 3,110 | 23.9 |
|  | ED Fig. 2d (right) | 5.49 | 22,469 | 3,544 | 17.8 |
| **Average** |  |  |  | **2,831** |  |

**Extended Data Fig 1.**
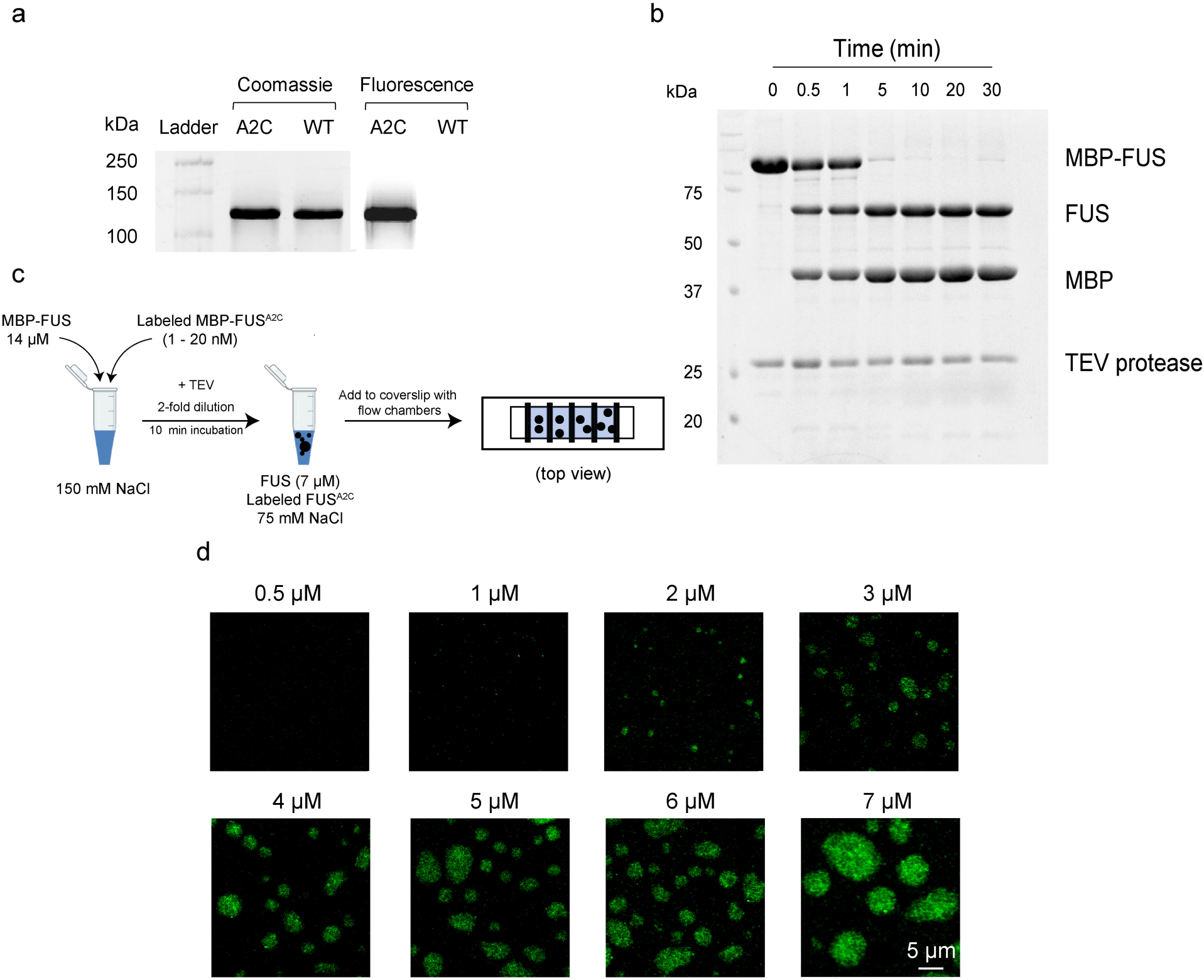
Preparation of FUS Condensates. **a,** Labeling of FUS^A2C^ with a maleimide dye. The labeling specificity of the A2C cysteine in FUS^A2C^ was determined using incubation of wildtype (WT) MBP-FUS and MBP-FUS^A2C^ with a 15-fold molar excess of Alexa647 maleimide (see Methods). The absence of fluorescent dye modification of MBP-FUS indicates that the four cysteine residues in the zinc finger domain (C428, C433, C444, C447) of FUS that are coordinated to Zn^2+[31]^ were protected from reaction with a maleimide dye. **b,** Kinetics of MBP removal by TEV protease activity. To remove the MBP and 6xHis tags from the FUS fusion protein (Fig. 1a), MBP-FUS (7 µM) was incubated with 3 units of TEV protease. Reactions were quenched after different time periods by addition of SDS-PAGE sample buffer and immediate heating to 100°C for 10 minutes. After 10 minutes, 99% of MBP-FUS was cleaved to yield separate MBP and FUS proteins. The MBP and TEV protease remained present in the resultant condensate mixture. **c,** Preparation of condensates for microscope observation. MBP-FUS (14 µM) and a low concentration of dye-labeled MBP-FUS were mixed (total volume of 5 µL) in a 150 mM salt buffer. The reaction mixture was diluted with an equal volume of no salt buffer containing 3 units of TEV protease. After 10 minutes, the mixture was added to flow chambers for imaging (see Methods). **d,** Estimation of *c*_sat_ for FUS. Different concentrations of MBP-FUS were mixed with ∼5 nM of MBP-FUS^A2C^-JF549. The presence of condensates was assessed ∼30 minutes after TEV protease addition by wide-field microscopy. From these images, *c*_sat_ ≈ 1 µM.

**Extended Data Fig 2.**
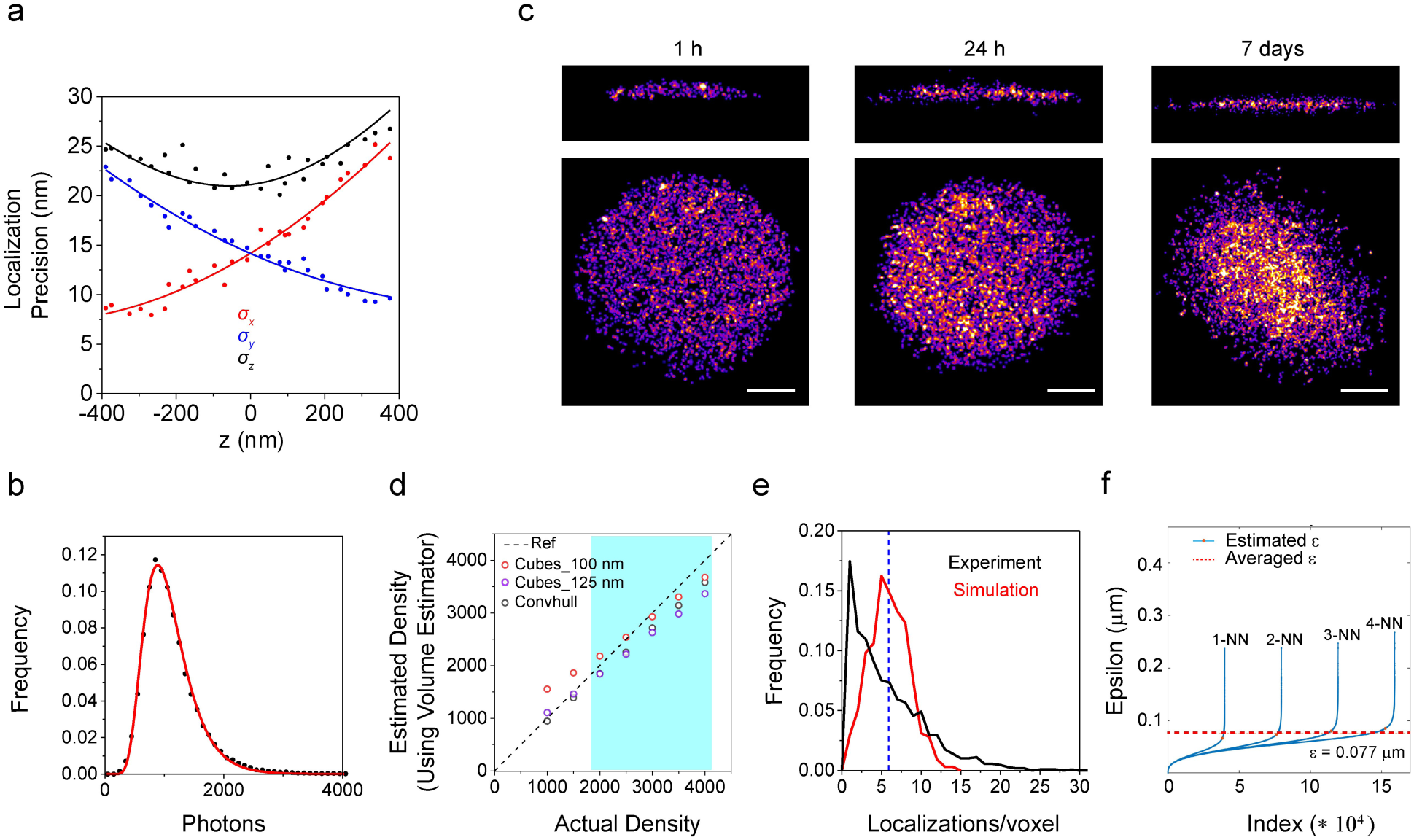
Generating 3D Localization Density Maps using Astigmatism Imaging. **a,** Localization precision. The *z*-dependent precisions in *x*, *y*, and *z* (α*_x_*, α*_y_*, and α*_z_*) for 60 nm root mean square deviation (RMSD) astigmatism were generated from repeated localizations of 0.1 µm fluorescent microspheres, as described earlier^36^. Precision values are normalized assuming 1000 photons/localization. **b,** Photon distribution histogram for FUS^A2C^-JF635b. Molecules were imaged at 30 ms/frame using 641 nm excitation (∼10 kW/cm^2^ measured at the sample). The mean of the log-normal fit is 1075 photons. **c,** PALM images. PALM images of FUS condensates containing 5-10 nM FUS^A2C^-JF635b were generated using the ThunderSTORM plugin in ImageJ. **d,** Comparing methods to estimate the condensate volume from simulated distributions. In the density range of the experiments (*shaded blue region*; see **Supplementary Table 2**), the cube volume estimator (see Methods) with a 100 nm cube side length performed better than 125 nm cubes and as well or better than MATLAB’s convexhull algorithm. **e,** Experimental and simulated densities. Frequency histograms are shown for experimental (FUS condensate aged for 7 days) and simulated (cylindrical volume) data with identical densities (3,169 localizations/µm^3^) for 100 nm cubic voxels from a 100 nm thick *xy* slice. The vertical dashed line indicates the average number of localizations (5.9 localizations/voxel). The experimental densities are more broadly distributed, indicative of non-randomness and consistent with clustering. **f,** The distance threshold (ε), or search radius, for DBSCAN analysis. Simulated data were used to estimate ε, which is defined as the knee of the *k*-nearest neighbors (kNN) curve. The knee is the point on the kNN curve closest to the point at which the curve’s asymptotes intersect. For the experimental densities observed, the average ε (= 0.077 µm) from 1NN to 4NN was used.

**Extended Data Fig 3.**
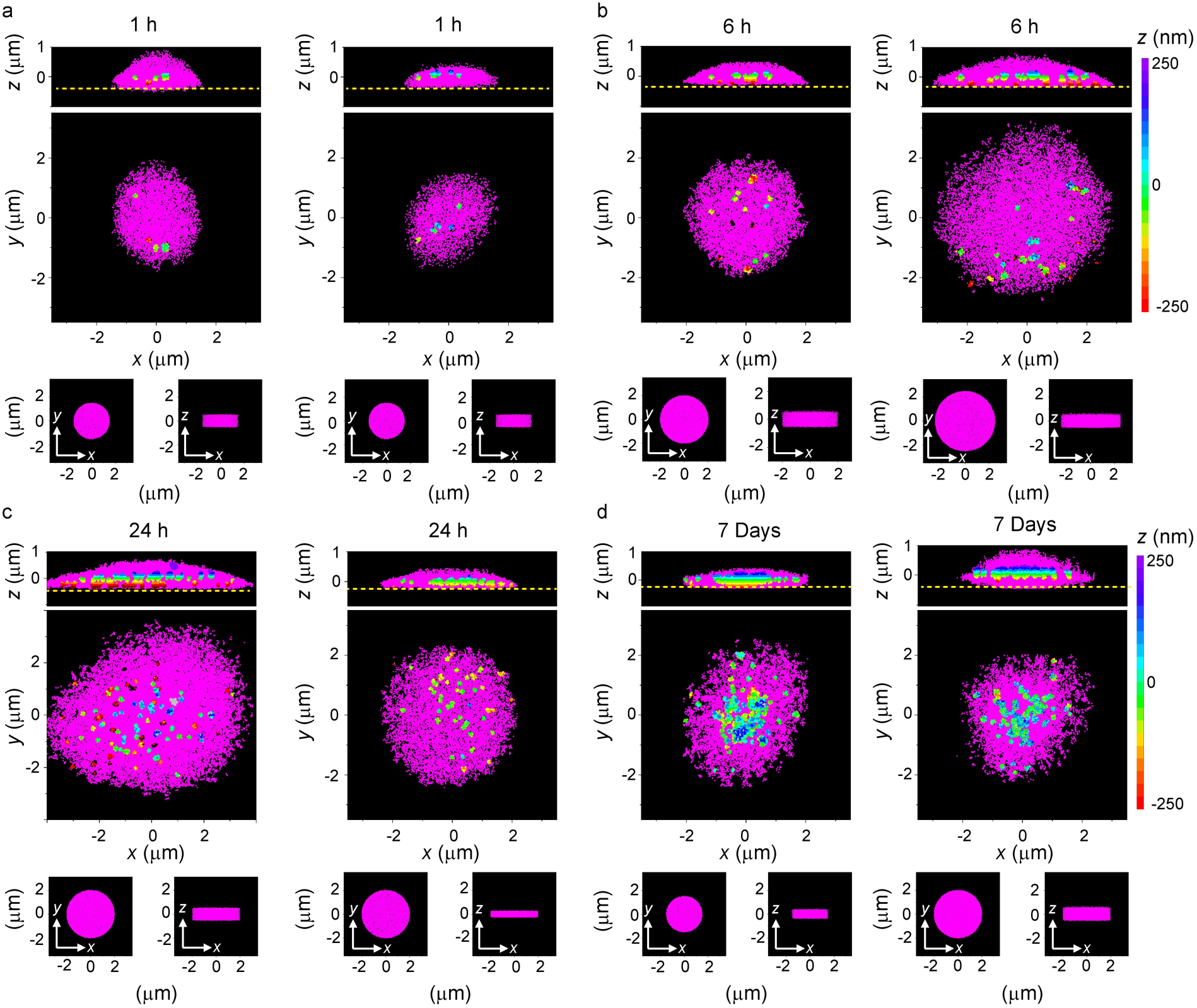
Increased Clustering with Longer Aging Times. **a-d,** 3D localization density maps of FUS condensates at four aging time points. These are additional replicates of the density maps from condensates shown in Fig. 1g**-l** that are analyzed and plotted in the same manner with experimental data on the top and simulated data on the bottom. In **a-d** (1 h, 6 h, 24 h, 7 days), two different condensates are shown, and all are distinct from Fig. 1. As in Fig. 1, there is a noticeable increase in clustering with longer aging times.

**Extended Data Fig 4.**
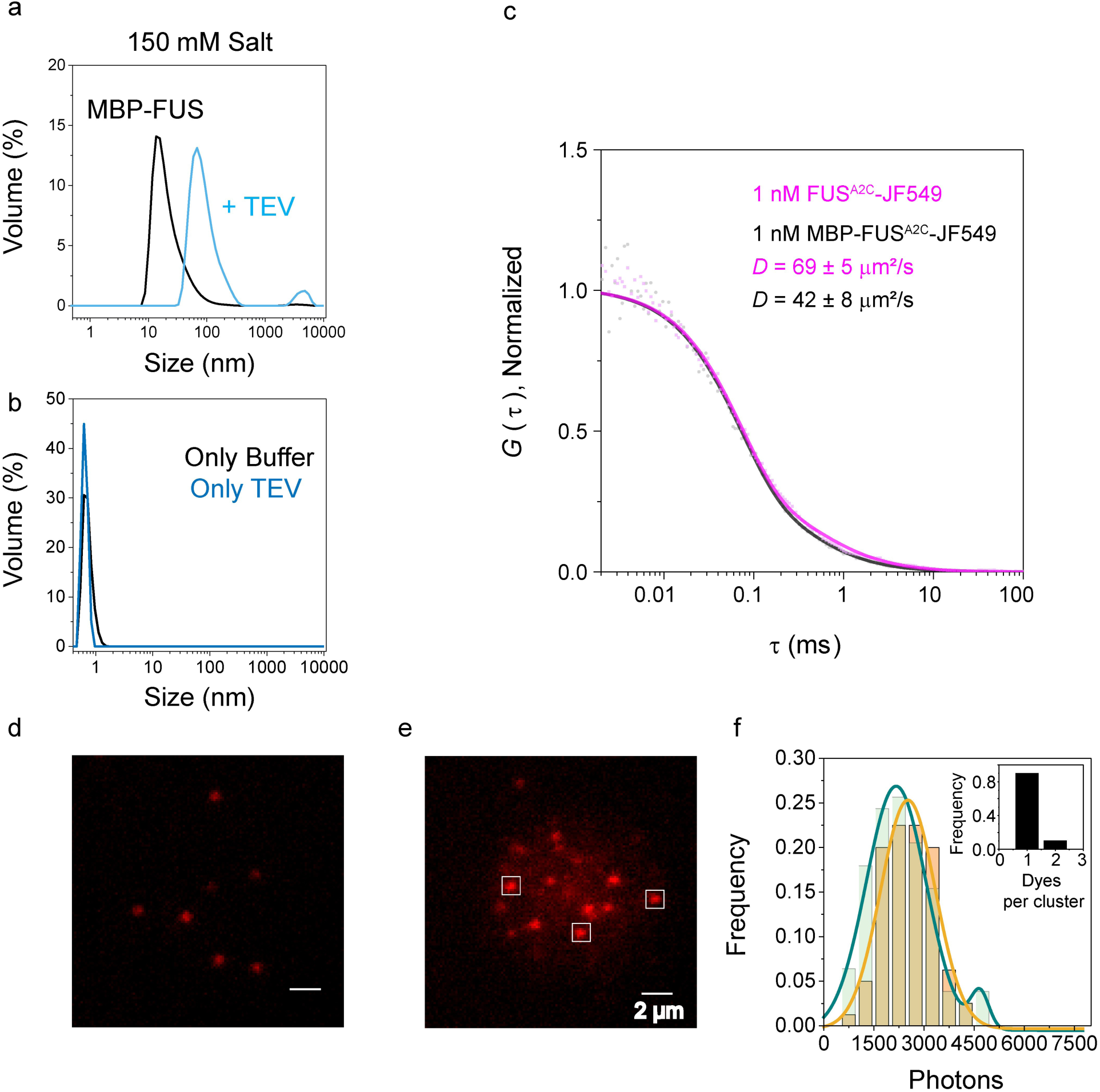
Size of FUS Assemblies within Condensates and at Low Concentration. **a,b,** DLS control experiments for nanoscale FUS assemblies at low concentration. DLS experiments performed under the same conditions of Fig. 2c except that the salt concentration was not diluted upon TEV addition (**a**, see **Extended Data Fig 1b**), only buffer was added (**b**; i.e., no MBP-FUS or TEV), or only TEV was added (**b**; i.e., no MBP-FUS). The solution in **a** initially contained 0.5 µM MBP-FUS (*black*) and was assessed again after TEV addition (*blue*). Note that the assemblies formed upon TEV addition are ∼100 nm in diameter, about half the size observed at 75 mM NaCl (Fig. 2c). **c,** Diffusion coefficients of MBP-FUS and FUS in 20 mM sodium phosphate, 2.5% glycerol, 200 mM NaCl, pH 7.5. A single component fit including a triplet state was used to determine diffusion coefficients (EX = 561 nm, 40 MHz pulse frequency, 20 µW, 300 s acquisition time). FUS^A2C^-JF549 was generated from MBP-FUS^A2C^-JF549 by adding TEV protease 10 minutes before beginning acquisition. *D* = 69 µm^2^/s for FUS^A2C^-JF549 indicates a hydrodynamic radius of ∼3.1 nm (estimated by the Stokes-Einstein-Sutherland equation). **d-f,** Number of dye molecules within condensate puncta. The intensity of MBP-FUS^A2C^-JF549 sparsely distributed on the coverslip surface (**d**) and FUS^A2C^-JF549 (1 nM) within FUS condensates aged for 1 h (**e**) were compared (**f**) under identical narrow-field acquisition conditions (100 ms/frame; EX = 561 nm, 8 kW/cm^2^). The number of photons per spot was determined within a 12×12 pixel region (see white boxes, 80 nm square pixels) corrected for the surrounding background. The intensity histogram (**f**; *green*, within condensate; *orange*, on surface) suggests that ∼90% of the puncta in the condensates have intensities corresponding to a single fluorophore. Considering a FUS:FUS^A2C^-JF549 ratio of 0.00014, these data suggest that the immobile puncta are an assembly with a maximum of ∼7000 molecules (corresponding to a diameter of ∼100 nm if close-packed, but a hydrated assembly is expected, suggesting a diameter in the 100-200 nm range). Note that the monomer control sample for these experiments (**d**) was MBP-FUS^A2C^-JF549 diluted from a 1 µM stock solution; since no large aggregates were observed upon dilution, which would yield significantly brighter intensity spots, the observed puncta in the condensates (**e**) were not insoluble aggregates present in the MBP-FUS stock solutions.

**Extended Data Fig 5.**
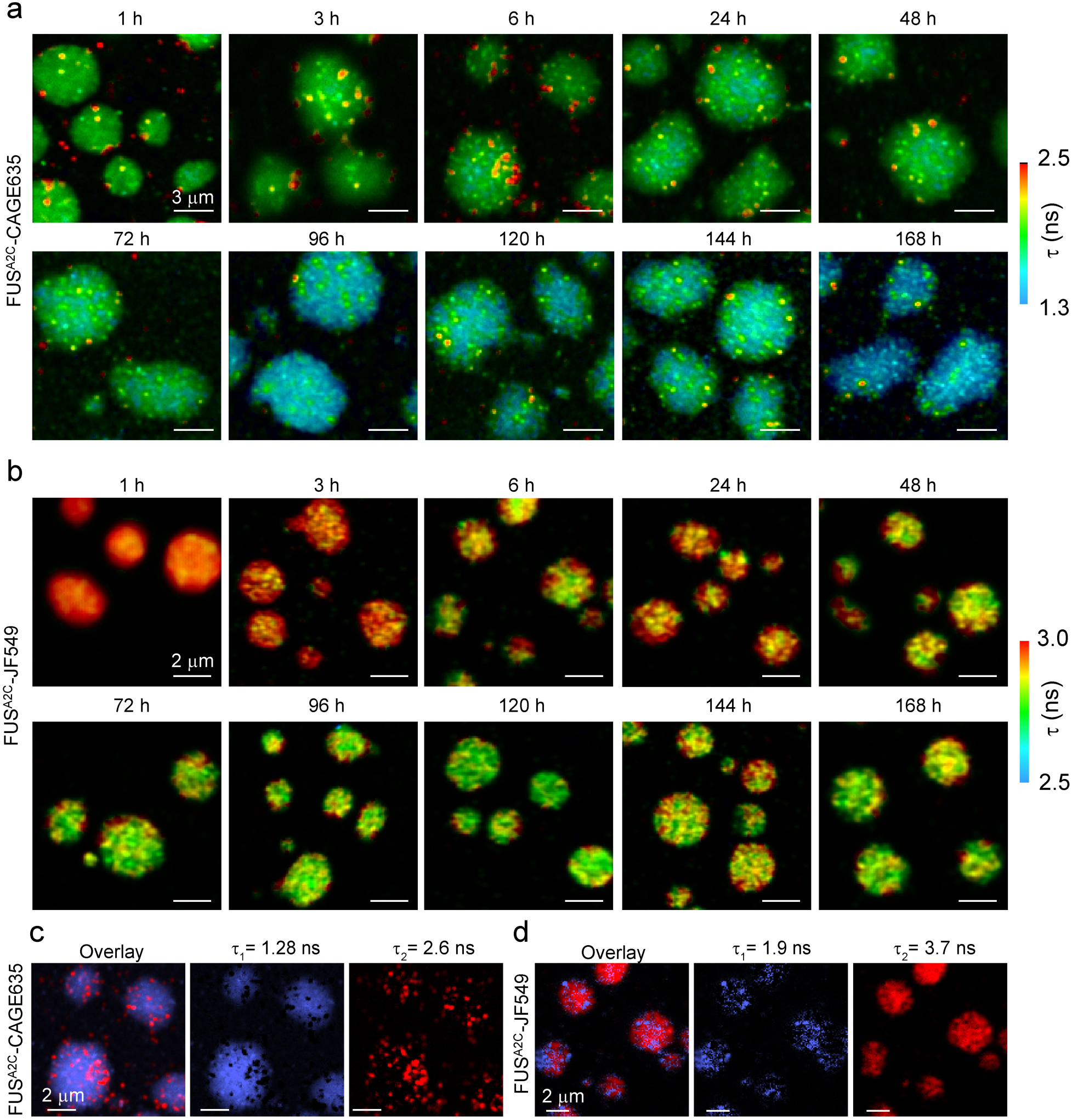
FLIM of FUS Condensates. **a,b,** FLIM images of condensates after various aging times. FUS (7 µM) condensates were formed using 10 nM FUS^A2C^-CAGE635 (**a**) or FUS^A2C^-JF549 (**b**). A double-exponential decay model was used to fit the photon arrival-time data for each condensate, revealing a decrease in the global fluorescence lifetime with increasing condensate age. The color scale represents the amplitude-weighted average fluorescence lifetime per pixel. CAGE635 reveals more pronounced puncta than JF549, and it displays a larger age-dependent average lifetime decrease, as highlighted in Fig. 2e. **c,d,** Separation of two lifetimes by pattern matching. The two lifetime components from the 6 h images in **a** and **b** were extracted from the fits and their amplitudes in each pixel were plotted as intensity for FUS^A2C^-CAGE635 (**c**) and FUS^A2C^-JF549 (**d**), respectively. The longer lifetime components, associated with clustered regions, are shown in *red*, while the shorter lifetime components, which are more uniformly distributed across the condensates, are shown in *blue*.

**Extended Data Fig 6.**
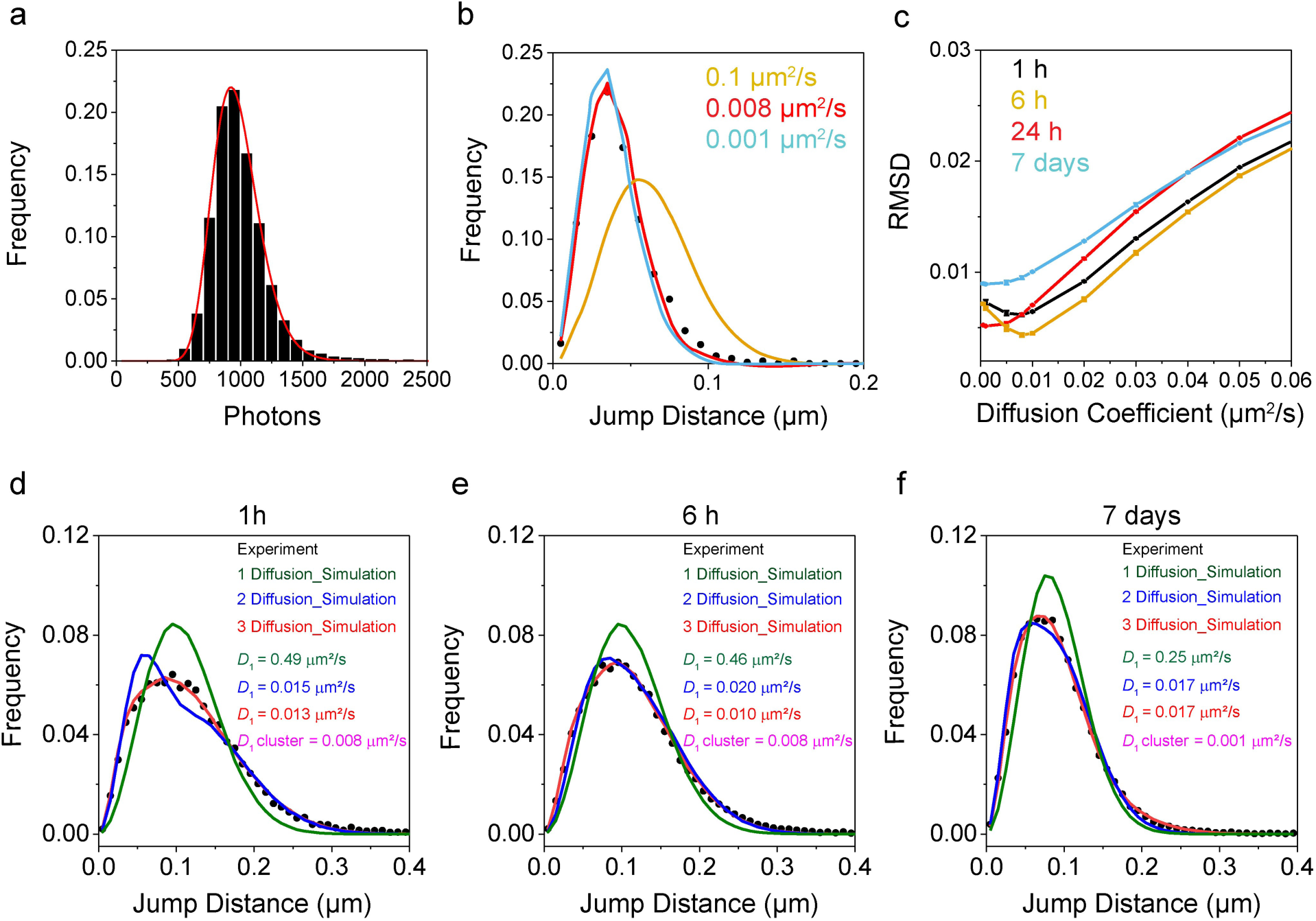
Diffusion Analysis of 3D Tracking Data from Astigmatism Imaging. **a,** Photon distribution histogram for FUS^A2C^-CAGE635. Tracking experiments were performed at 5 ms/frame for ∼25 min as in Fig. 3a and yielded an average photon count of 971 photons (16,103 localizations; 24 h aging time). All localizations within *z* = 0±250 nm and ≥ 500 photons/localization were used for all analyses, which yielded average experimental precisions of σ_x_ = 12.7 nm, σ_y_ = 15 nm and σ_z_ = 21 nm, estimated from the *z*-dependent precision curves in **Extended Data Fig. 2a**. **b,** Diffusion coefficient of the confined fraction. Confined tracks obtained after 6 h of aging were isolated and the jump step histogram was compared with a simulated distribution assuming a single diffusion coefficient (**Eq. 1**) and including the experimental precision (**Jump Step Histogram Fitter**). These data are the same as that in Fig. 3b; the simulated distributions shown here assume the diffusion constant as indicated by the color. At lower values of *D*, the precision dominates the observed jump distribution, and the curves are difficult to distinguish for different values of *D*. **c,** Determining the best fit. The root mean square deviation (RMSD) between experiment and simulated curves for different *D* values was used to identify the best fit. Here, the best fit for the 6 h time point occurred at ∼0.008 µm^2^/s (see fit in **b**). **d-f,** Diffusion coefficients for the total population. Having determined a diffusion coefficient for the confined fraction (**b,c**), the jump step histograms for entire trajectory datasets at the different timepoints were then generated and fit with simulated data (minimizing the RMSD) assuming one, two, or three diffusion coefficients (**Eq. 3**) and including the experimental precision (**Jump Step Histogram Fitter**). While two diffusion coefficients appeared sufficient for a reasonable fit in some cases, visual inspection and the RMSD values indicated that the three diffusion coefficient model yielded the best fits. The color of the *D*_1_ coefficients in the figure panels indicate how many diffusion coefficients were assumed for the fitting.

**Extended Data Fig 7.**
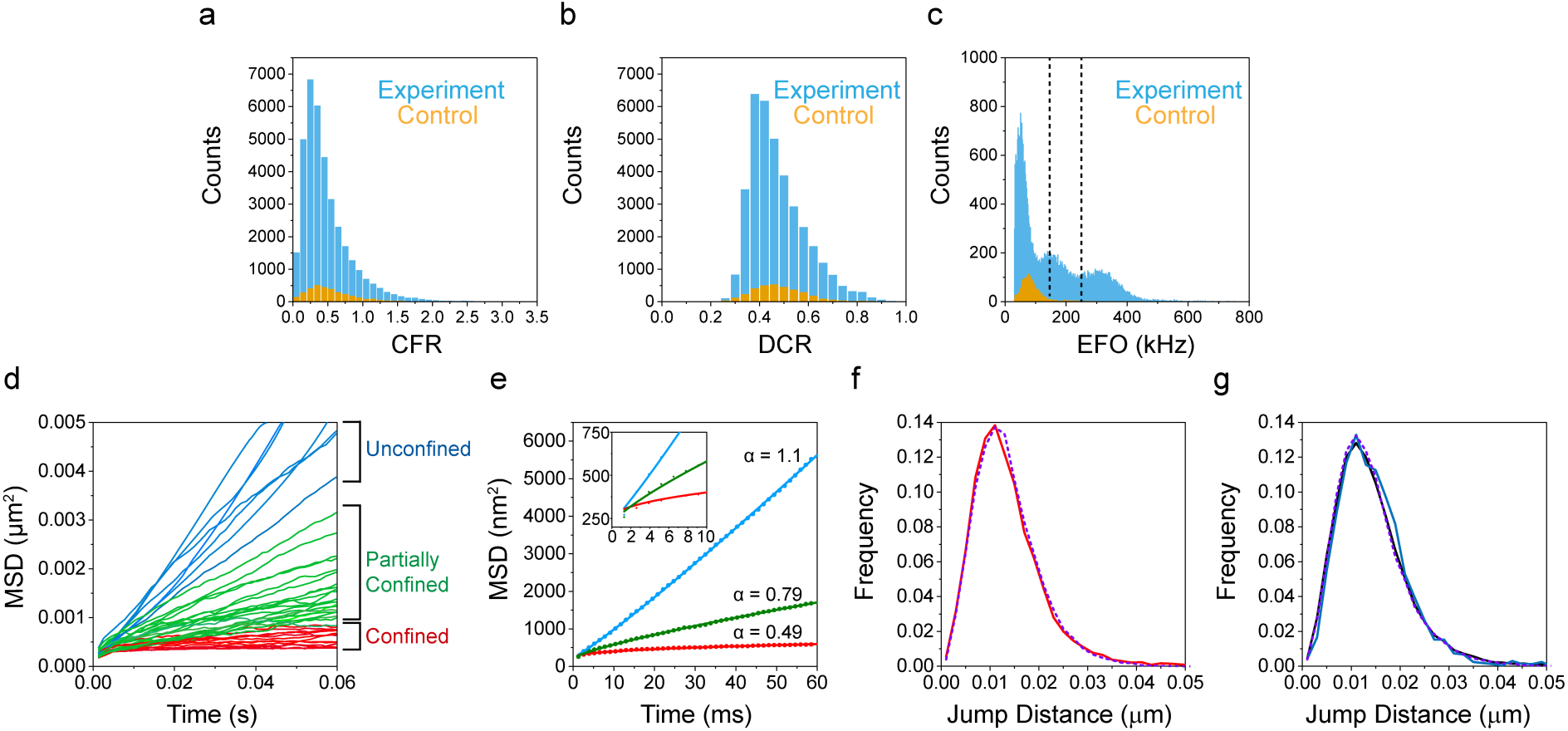
Diffusion Analysis of 3D Tracking Data from MINFLUX Imaging. **a-c,** Post-acquisition filtering of the MINFLUX data. The center frequency ratio (CFR), detector channel ratio (DCR) and the emission frequency at offset (EFO), and what they mean and how they are used was described earlier^52^. Control data was acquired by imaging FUS condensates without FUS^A2C^-CAGE635 and is considered the background noise from the experiment. The CFR (**a**) and DCR (**b**) distributions obtained over 60 minutes were virtually indistinguishable in shape between control (*orange*) and experimental (*blue*) conditions, yet the control observations were substantially lower in number (∼10% of experimental). In the experimental EFO (**c**) histogram, the first peak is considered noise, and the second and third peaks are considered to arise from one and two dyes, respectively. Thus, for initial screening, tracks with an average EFO between 150 kHz and 250 kHz (dashed vertical lines) were selected. While the control CFR and DCR histograms indicate individual fluorophores within the background that have a similar emission spectrum to CAGE635, the EFO histogram indicates that the emission intensity is lower. MINFLUX localizations are typically considered to be poor for a CFR > 0.8^[52]^. Therefore, tracks were retained only if their average CFR value was < 0.8 and if the track contained < 10% of localizations with CFR > 0.8. **d,e,** Confinement within FUS^A2C^-CAGE635 tracks. Tracks with > 150 localizations (45 of 87) were analyzed via mean squared displacement (MSD) analysis (two tracks were eliminated due to large fluctuations in the MSD trajectory). Variable levels of confinement were identified, as indicated (**d**). Note that a maximum of ∼30% of the MSD trajectories are shown^58^. Tracks identified as confined (α < 0.6), partially confined (0.6 < α < 0.9), and unconfined (0.9 < α), were averaged and fit with **Eq. 5** to obtain *D* = 0.010 µm^2^/s, the indicated α values, and σ = 10-11 nm (**e**). **f,g,** Simulated jump step histograms. The jump step histograms from Fig. 4c were approximated with jump step histograms simulated by **Trajectory Simulator** (see Methods). The confined (**f**), unconfined and all (**g**) tracks (same colors as in Fig. 4c) were simulated (*purple dashed*) with the same parameters [two diffusion coefficients of 0.008 µm^2^/s (75%) and 0.05 µm^2^/s (25%) and α*_x_* = α*_y_* = 1.608α*_z_* = 4.8 nm], except that the level of confinement to a spherical volume varied (*r* = 25 and 50 nm, respectively).

**Extended Data Fig 8.**
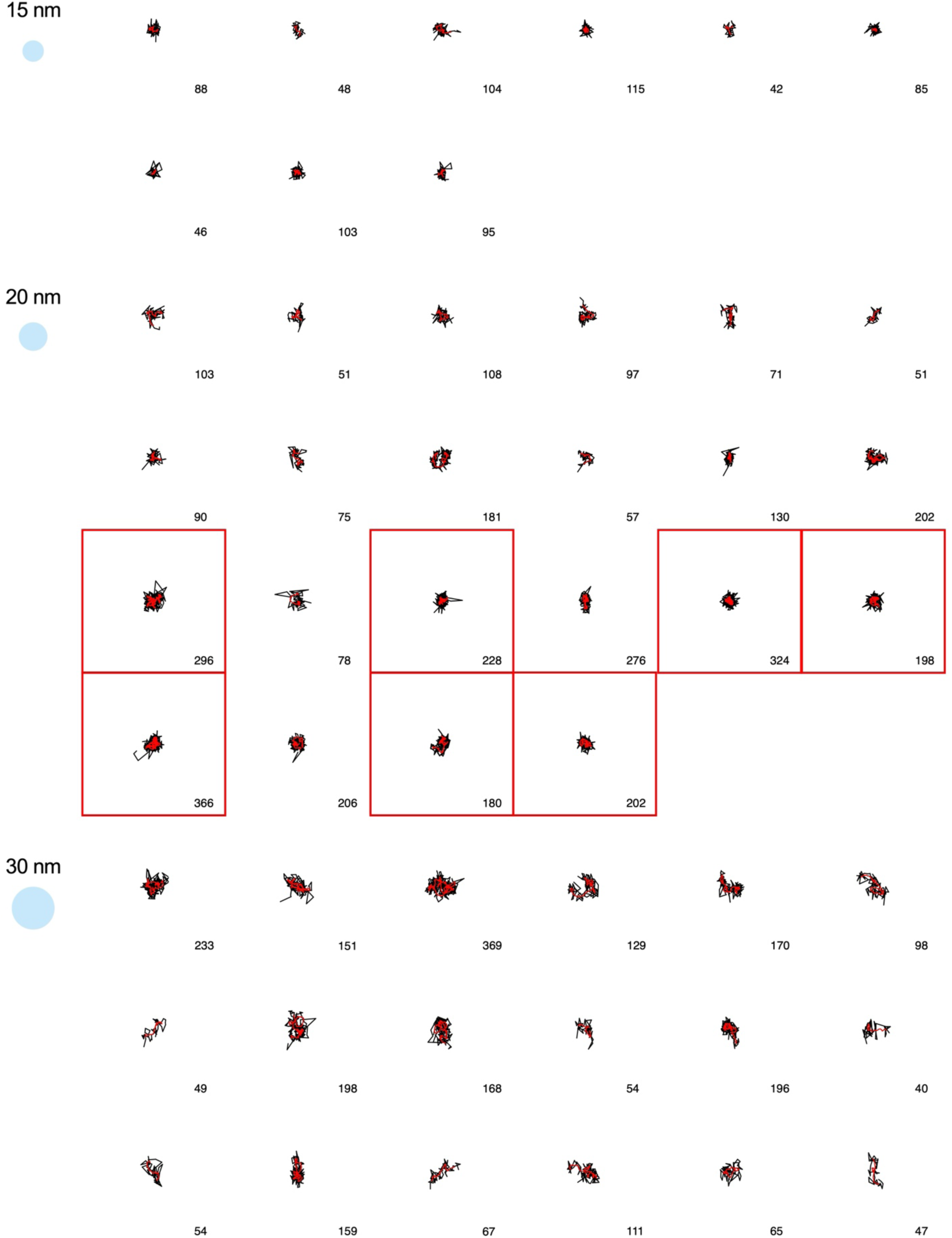
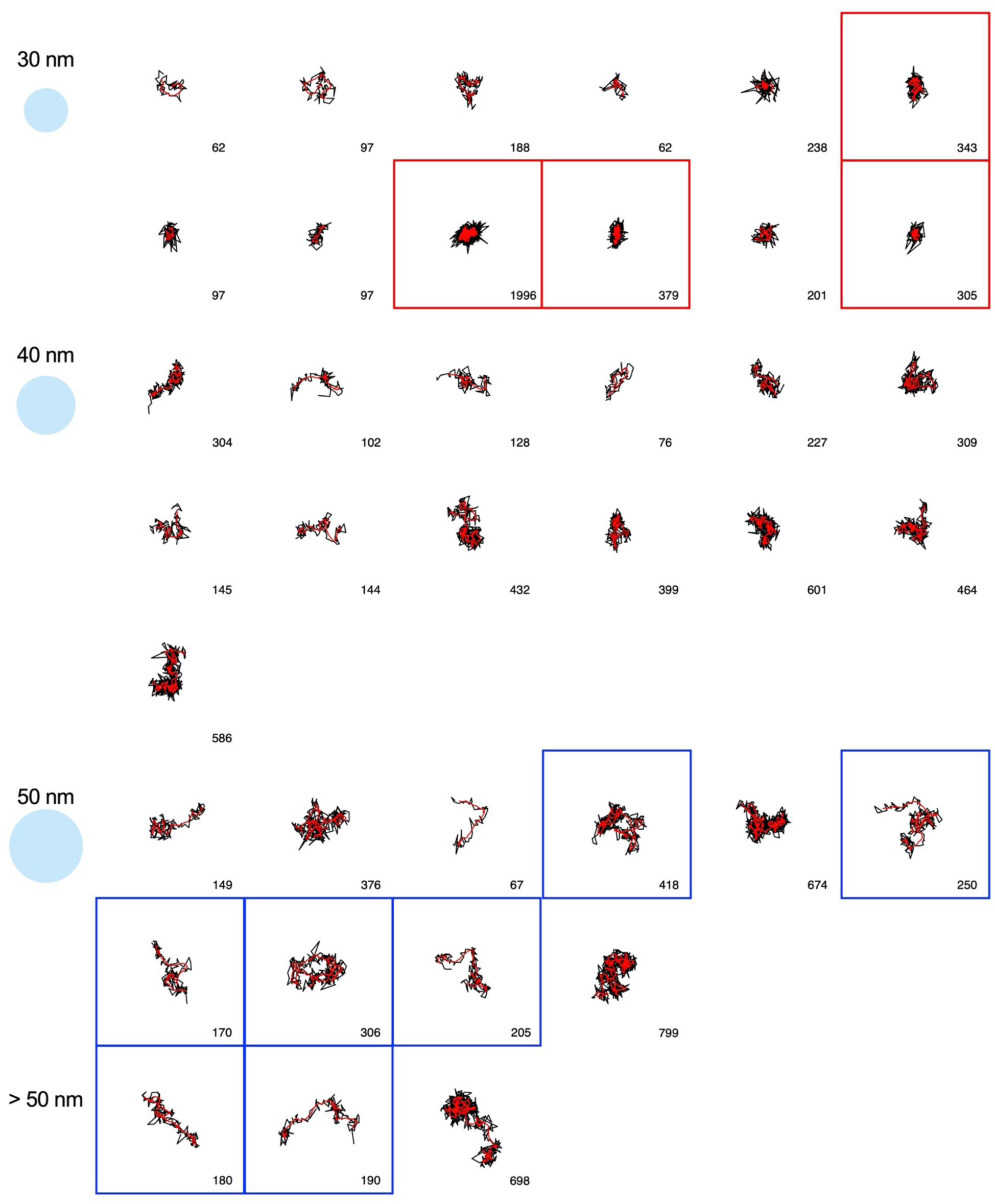
MINFLUX tracks. All 87 MINFLUX tracks that survived CFO and EFO filtering (see **Extended Data Fig. 7a-c**) are shown organized in groups defined by the approximate range of motion (*blue disks*). The number of points in each trajectory is indicated at the bottom right of each panel. The trajectories (*black*) are shown with a moving centroid calculated from 10 localizations (*red*). Trajectories boxed in *red* and *blue* were used to generate the confined and unconfined jump step histograms in Fig. 4c, respectively.

**Extended Data Fig 9.**
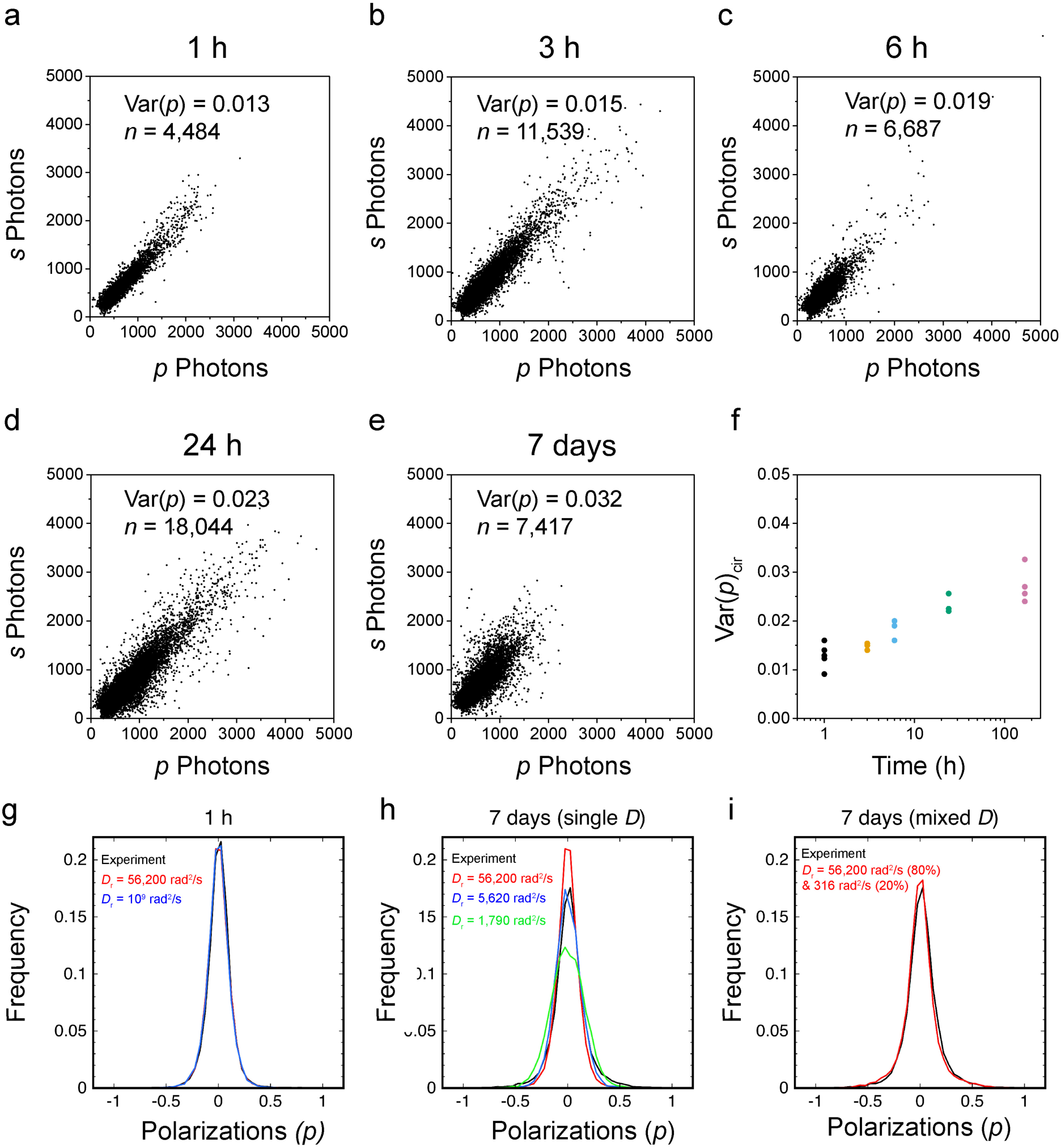
Photon Scatterplots, Individual Var(*p*)_cir_ Measurements, and Simulated Polarization Histograms for SiMRoD Measurements. **a-e,** Photon scatterplots. Sample distributions of the photons collected in the *p-* and *s-*polarization channels from single condensates after different aging times. Each dot represents the photons collected in 5 ms for one FUS^A2C^-JF635b molecule within a condensate. A minimum of 200 photons/localization in at least one of the channels was required for inclusion in Var(*p*) calculations. An increase in scatter relative to the diagonal indicates an increase in Var(*p*). **f,** Individual Var(*p*) values of all of the condensates analyzed. These are the raw values for Fig 5j. **g-i,** Simulated polarization histograms. Experimental SiMRoD data (*black*) after 1 h of aging time were well approximated by simulated data (**g**) in which a single *D_r_* was assumed [56,200 rad^2^/s or 10^9^ rad^2^/s, Var(*p*)_cir_ = 0.012 for both]. In contrast, data after 7 days of aging time were not well approximated by the same simulation data or any single *D_r_* distribution (**h**). Instead, two rotational diffusion coefficients [*D_r_*_(1)_ = 56,200 rad^2^/s (80%), *D_r_*_(2)_ = 316 rad^2^/s (20%), Var(*p*)_cir_ = 0.027] were required to best approximate the experimental distribution (**i**).

## Supplementary Video Captions

**Supplementary Video 1.**

**Imaging of FUS^A2C^-JF635b.** This video shows the self-blinking behavior of the dye JF635b. FUS (7 µM) was spiked with ∼10 nM FUS^A2C^-JF635b and imaged using astigmatism microscopy at 30 ms/frame for ∼20 min.

**Supplementary Video 2.**

**Narrow-field imaging of FUS Condensates aged for 1 h.** This video shows the slow movement of FUS^A2C^-JF549 (∼0.5 nM) within a FUS condensate aged for ∼1 h and imaged using narrow-field epifluorescence at 200 ms/frame. Two condensates merge during this video.

**Supplementary Video 3.**

**Narrow-field imaging of FUS Condensates aged for 6 h.** This video shows the slow movement of FUS^A2C^-JF549 (∼0.5 nM) within a FUS condensate aged for ∼6 h and imaged using narrow-field epifluorescence at 200 ms/frame.

**Supplementary Video 4.**

**Narrow-field imaging of FUS Condensates aged for 7 days.** This video shows the slow movement of FUS^A2C^-JF549 (∼0.5 nM) within a FUS condensate aged for ∼7 days and imaged using narrow-field epifluorescence at 200 ms/frame.

